# APC coordinates GSK3 phosphorylation of SETD8 to suppress colorectal cancer

**DOI:** 10.1101/2024.10.03.616482

**Authors:** Zvi Cramer, Keara Monaghan, Ricardo Petroni, Xin Wang, Stephanie Adams-Tzivelekidis, Kayla Durning, Melissa S. Kim, Yuhua Tian, Nicolette M. Johnson, Nicolae A. Leu, Simone Sidoli, Ning O. Li, M. Andres Blanco, Christopher J. Lengner

## Abstract

Colorectal cancer (CRC) is the second-leading cause of cancer-related deaths with increasing incidence globally. Mutations in the tumor suppressor APC initiate CRC at least in part by preventing the GSK3 kinase from phosphorylating β-CATENIN, leading to its constitutive stabilization and transactivation of mitogenic target genes. While the importance of β-CATENIN phosphorylation by GSK3 is well-established, APC regulation of GSK3 activity upon other targets with potential oncogenic relevance are not understood. Here, we identify the H4K20 methyltransferase SETD8 as target of APC-coordinated GSK3 phosphorylation in the intestinal epithelium. We found that phosphorylation by GSK3 restrains the oncogenic activity of SETD8, with loss of phosphorylation sensitizing mice to oncogenic insults. Mechanistically, phosphorylation alters the role of SETD8 in transcriptional regulation, most notably by preventing it from activating oncogenic YAP signaling and a fetal-like transcriptional program. These results underscore the importance of SETD8 in CRC and represent a novel β-CATENIN -independent oncogenic consequence of APC loss.

**Significance:** GSK3 is thought to restrain colorectal cancer primarily by phosphorylation of β-CATENIN. We show that GSK3 also phosphorylates SETD8, preventing SETD8 activation of oncogenic programs including YAP-driven fetal-like gene expression.

## Introduction

As the second-leading cause of cancer-related deaths in the United States, colorectal cancer (CRC) is in desperate need of therapeutic innovation^1^. Mutation of the *APC* tumor suppressor gene is a recurrent driver of CRC tumorigenesis, occurring in ∼80% of CRC patients^2^. In normal colorectal epithelium, APC is a critical component of the ‘destruction complex’, which contains the scaffolding protein AXIN, glycogen synthase kinase 3 (GSK3), casein kinase 1 (CK1), and the transcriptional effector of the canonical WNT signaling pathway, β-CATENIN^3^. In epithelia, most β-CATENIN is associated with adherens junctions, with cytosolic/nuclear pools under tight regulatory control^3^. In the absence of external WNT ligands, APC and AXIN coordinate the phosphorylation of cytosolic β-CATENIN by GSK3 and CK1, leading to recognition and degradation of β-CATENIN by the ubiquitin-proteasome system^4^. Upon WNT ligand-receptor interaction, APC is released from the destruction complex, leading to β-CATENIN stabilization and translocation into the nucleus. There, β-CATENIN interacts with T-cell factor/lymphoid enhancer factor (TCF/LEF) transcription factors to promote the expression of genes supporting mitotic self-renewal of adult stem cells in the intestinal/colonic epithelium^3,6^.

In CRC, *APC* mutations are overwhelmingly truncating mutations in one allele that remove all AXIN binding sites (SAMP repeats) and some β-CATENIN binding sites and occur in conjunction with complete loss of the second allele^7^. Thus, the field has generally accepted that truncation of APC drives CRC tumorigenesis primarily by aberrant activation of β-CATENIN transcriptional activity. Indeed, the β-CATENIN target gene *Myc* is necessary for tumorigenesis induced by *Apc* loss^8^. Nevertheless, the overrepresentation of APC loss of function mutations relative to activating mutations in other components of the canonical WNT/β-CATENIN pathway suggests that APC may have additional tumor suppressive activities^2^. Indeed, mounting evidence has implicated APC in other cellular pathways^5,7^. For example, APC plays roles in DNA-damage repair, microtubule organization and cell adhesion^9–11^. Furthermore, Valvezan et al. recently demonstrated that APC potentiates GSK3 phosphorylation of other substrates besides β- CATENIN, including Glycogen Synthase, Tau, and components of the mTOR pathway^5^. However, the pervasiveness and importance GSK3 targets beyond B-CATENIN in APC-mutant CRC is unknown.

Here, we test the hypothesis that loss of APC disrupts GSK3 activity on targets beyond β- CATENIN, and that this dysregulation impacts CRC tumorigenesis. We systematically define phosphosites lost upon acute conditional deletion of either GSK3 or APC in murine intestinal epithelium. We identify SETD8 Thr138 as a phosphorylation event catalyzed by GSK3 that is dependent on APC. SETD8 is a member of the SET-domain family of methyltransferases and is the only known catalyst of histone H4K20 to H4K20me1, a post-translational modification that plays roles in cell cycle progression, DNA repair, and transcription^12^. SETD8 also has non-histone substrates, including P53 and CD147^13,14^. SETD8 has also been implicated in human cancer. For example, a polymorphism found within the 3’ untranslated region of SETD8 that is predicted to disrupt the binding of miR-502 is associated with elevated SETD8 expression and increased cancer risk in early-onset breast cancer, clear cell renal carcinoma and CRC^15–17^. Moreover, SETD8 is an essential gene in breast cancer, CRC, lung cancer and medulloblastoma^13,18–20^. Thus, accumulating evidence suggests SETD8 may represent a therapeutic target in a variety of cancer types.

We found SETD8 Thr138 phosphorylation to be tumor-suppressive in tumor organoids and *in vivo* mouse models of CRC, where it suppresses intestinal fetal-like and cholesterol biosynthesis gene expression programs. This work represents a shift in the dogmatic model of APC in CRC and is likely to identify novel points for therapeutic intervention in this widespread and deadly disease.

## Materials and Methods

### Phospho-mass spectrometry of murine intestine

Intestinal crypts were harvested by scraping and EDTA incubation and 0.8mg protein from each sample was digested in-solution with trypsin and cleaned up with SepPak C18. Phosphopeptide enrichment was performed using Titansphere Phos-TiO column (GL Sciences Inc.) with two rounds of sequential enrichment. The eluted phosphopeptides were combined and analyzed by LC-MS/MS on a Q Exactive HF mass spectrometer using an extended 4-hour LC gradient. MS/MS spectra generated from the LC-MS/MS runs were searched against the UniProt mouse database (www.uniprot.org; 10/01/2018) using the MaxQuant 1.6.2.3 program with full tryptic specificity. The “Match between runs” feature was used to transfer identifications across experiments to minimize missing values. Proteins and peptides with S, T, Y phosphorylation were required to have a minimum score of 40 and delta score of 6. The false discovery rates for protein, peptide, and site identifications were set at 1%. The intensity values were normalized by dividing against median value. Missing values (Intensity = 0) were imputed with a small value derived from the minimum value of the dataset divided by 2. Averages were calculated for each group based on the imputed normalized intensity and fold changes between two groups were determined from these average values.

### Genotyping validation of *Apc*^flox/flox^ and *Gsk3*α/β^flox/flox^ recombination

Genotyping for *Apc*^flox/flox^ was performed using previously described primers^58^ while *Gsk3*α/β^flox/flox^ was performed using custom-designed primers and previously described primers^23^. Both PCR reactions were carried out using mouse DNA isolated by DNA Micro Kit (Qiagen #56304) using Ex Taq (Takara #RR001A).

The following primers were used:

> *Apc*-P3: 5’-GTTCTGTATCATGGAAAGATAGGTGGTC-3’, *Apc*-P4: 5’- CACTCAAAACGCTTTTGAGGGTTGATTC-3’; *Apc*-P5: 5’- GAGTACGGGGTCTCTGTCTCAGTGAA-3’; *Gsk3α*-F1: 5’-TGTACCTGCCCTGCAAAAGG-3’; *Gsk3α*-R1: 5’-GCCTGTAACCTCACCAAACAC-3’; *Gsk3α*-R2: 5’- CTGTGGTAGAATAACTGTCTGG-3’; *Gsk3β*-F1: 5’- CCTCATGTGACTATATTCTCAAAGG-3’; *Gsk3β*-R1: 5’- GATAGCTCAGTAGGTCAAAGGG-3’; *Gsk3β*-R2: 5’- CTTGTCCCTGCAATCAAAGC-3’

### Cell culture

HEK293T cells were cultured in DMEM (Life Technologies #11965-084) supplemented with 10% Fetalplex (Gemini #100-602), 100µM sodium pyruvate (Gibco #11360070), 1X non-essential amino acids (Gibco # 11140050), 1X penicillin–streptomycin (Gibco #15140122), and 55µM 2- mercaptoethanol (Gibco #21985023).

Tumoroids were cultured as previously described^59^. Briefly, organoids were maintained in complete medium composed of Advanced DMEM/F-12 (Thermo Fisher Scientific #12634010) supplemented with 1X penicillin–streptomycin (Gibco #15140122), 10μg/mL gentamicin (Gibco #15710064), 10mM HEPES (Thermo Fisher Scientific #15630-080), 1X B27 (Gibco #17504-044), 1X N2 (Invitrogen #17502-048), 1X GlutaMAX (Thermo Fisher Scientific #35050061), 1mM N-acetyl-cysteine (Sigma-Aldrich #A9165) and 1x conditioned media (derived from L-WRN cells). Organoids were passaged using Dispase (STEMCELL Technologies #07923) and Accutase (Sigma-Aldrich #A6964). Cells were seeded in 3:1 Matrigel:PBS and complete medium supplemented with 10µM Y-27632 (Selleck Chemicals #S1049). Monolayers were passaged as previously described^59^, with some modification. Monolayer cultures were passaged with 500U/mL collagenase IV (Gibco #17104019) and washed in 1mL of 0.5mM EDTA-PBS, before seeding on top of collagen I hydrogels (Corning #354249) in complete medium supplemented with 500 nmol/L A83-01 (Tocris #2939) and 10µM Y-27632.

All cells were routinely tested for Mycoplasma contamination.

### Generation of cDNA vectors and transient transfection

For stable expression of SETD8 in *CAPK* tumoroids, pUltra-Chili (a gift from Malcom Moore, Addgene #48687) was digested with XbaI (NEB #R0145) and BamHI-HF (NEB #R3136) and WT SETD8 or T140A/D SETD8 cDNA were ligated using T4 ligase (NEB #M0202). T140A/D SETD8 cDNAs were generated using the QuikChange Lightning Site-Directed Mutagenesis Kit according to the manufacturer’s instructions (Agilent #210518).

Human SETD8 cDNA was cloned with Q5 polymerase (NEB #M0493S) using the following primers:

> *SETD8*-cDNA-F: 5’-CGCTCTAGAATGGCTAGAGGCAGGAAGATGTGC-3’; *SETD8*-cDNA-R: 5’-CGCGGATCCCCACGTGGTTAGTGCTTCAGCCA-3’

Targeted mutagenesis was performed using the following primers:

> *SETD8-T140A*-F:5’-TGAAGCAGCAGAACCTCCAAAAGCTCCACCCTCAT-3’;*SETD8-T140A*-R: 5’-ATGAGGGTGGAGCTTTTGGAGGTTCTGCTGCTTCA-3’; *SETD8-T140D*-F: 5’-AGATGAGGGTGGATCTTTTGGAGGTTCTGCTGCTTCAGAT-3’;*SETD8-T140D*-R: 5’- CACAAGATGAGGGTGGCTCTTTTGGAGGTTCTGCTGCTTCAGATTTTTGG-3’

For transient expression of SETD8, 293T cells were transfected using PEI (PolyScience #24765- 1) for 5 hours and collected 48h post-transfection.

### Western blot analysis

Tumoroids or intestinal crypts were collected and lysed with RIPA Buffer (Abcam ab156034) supplemented with a protease/phosphatase inhibitor cocktail (Cell Signaling Technology #5872S), sonicated and spun down for 15 min at 14000g at 4°C. Protein concentration of resulting supernatant was determined the Pierce™ BCA Protein Assay Kit (Thermo Scientific #23225). Equal amounts of protein were resolved on a 10% SDS–PAGE gel and after transfer, membranes were blocked with 5% bovine serum albumin (BSA, Sigma-Aldrich #A7906) for 1 hour at room temperature and then incubated with indicated primary antibodies diluted in 5% BSA TBST overnight at 4°C. Membranes were then incubated with HRP conjugated secondary antibodies, which included secondary anti-rabbit antibody (CST#7074S, 1:1,000) or anti-mouse antibody (CST#7076S, 1:2000). Blots were visualized with SuperSignal West Pico PLUS Chemiluminescent Substrate (Thermo Scientific, #34577) and Bio-RAD Chemidoc TMMP imaging system. The following primary antibodies were used: anti-β-ACTIN (Abcam #ab6276, 1:5000), anti-SETD8 (CST #2996, 1:1000), anti-H4K20me1 (Abcam #ab9051), anti-histone H3, (Abcam, ab1791), and anti-histone H4 (CST #2935).

### Recombinant protein purification in bacteria

Rosetta bacteria (Millipore #70954-4) were transformed with pGEX-4T1 containing SETD8 WT, T140A or T140D cDNAs and grown until A600 reach 0.5-0.8. 1mM IPTG (Invitrogen) was then added and bacteria were grown overnight at 18-20⁰C with shaking. The next day, bacteria were lysed in protein lysis buffer (50mM Tris (pH 7.3), 250mM NaCl, 0.05% Triton X-100, 3mM 2- mercaptoethanol) supplemented with 0.5mM PMSF and sonicated at 50% power for 10 minutes (10 seconds on, 15 seconds off). Lysate was spun and supernatant was incubated with glutathione-sepharose beads (Sigma-Aldrich GE17-0756-01) overnight at 4⁰C. Beads were then washed six times with protein lysis buffer and eluted at 4⁰C for 15 minutes with protein lysis buffer supplemented with 10mM Glutathione and adjusted to pH 8. Elutant was then dialyzed overnight at 4⁰C with TAP-Wash buffer 50mM Tris (pH 7.9), 100mM KCl, 5mM MgCl_2_, 0.2mM EDTA, 0.1% NP-40, 10% glycerol, 2mM DTT). Protein concentration was quantified using the Pierce™ BCA Protein Assay Kit (Thermo Scientific #23225) according to the manufacturer’s instructions.

### Recombinant protein purification in insect cells

Sequence-confirmed pFastBac transfer vector containing sequences for APC-SAMP fragments^5^ were transformed into DH10Bac cells. Proper recombination of the transfer vector into the baculovirus genome was confirmed by PCR and positive bacmid DNAs were transfected into Sf9 cells and passage 1 (P1) virus stocks were recovered 120 hours post-transfection. A high-titer P2 virus stock was generated by infecting Sf9 at an MOI of ∼0.1, followed by incubation for 120 hours. For productions, 1 × 10^6^ Sf9 cells/ml in Sf900-III medium (Invitrogen) were infected with virus at an MOI of 1 and harvested 48 hours post-infection^60^.

### Radioactive kinase assay

500 ng of purified recombinant GST-SETD8 proteins were incubated with 3μL of recombinant GSK3α (Abcam ab42597) or GSKβ (Abcam ab63193) in kinase assay buffer (20mM Tris (pH 7.5), 10mM MgCl2, 5mM DTT) with or without 10μCi of ATP-γ-32P (PerkinElmer NEG002A250UC). Reactions were incubated at 30⁰C for 30 minutes and terminated with loading dye (Biorad #161074). Samples were run on an SDS-Page gel, stained/fixed with Imperial Protein Stain (Thermo Fischer Scientific #24615) and visualized with autoradiograph film (LabScientific).

### Methyltransferase assay

10-100nM of recombinant GST-SETD8 protein were incubated with 0.5μg of recombinant mononucleosomes (H3.1, Active Motif #81070) in methyltransferase buffer (30mM Tris (pH 8.0), 25mM NaCl, 0.5mM DTT) with or without 8μM S-adenosylmethionine (SAM) for 15 minutes at room temperature. Reactions were stopped by adding loading dye (Biorad #161074).

### Nucleosome binding assay

Nucleosome binding assays were performed as previously described^61^. 1.5μg of recombinant GST-SETD8 proteins were incubated with 1μg of recombinant mononucleosomes (H3.1, Active Motif #81070) and glutathione-sepharose beads (Sigma-Aldrich GE17-0756-01) in GRB buffer (20 mM HEPES pH 7.5, 10% glycerol, 25 mM KCl, 0.1 mM EDTA, 1 mM dithiothreitol (DTT), 2 mM MgCl2, 0.2% NP40) supplemented with 8μM S-adenosylmethionine (SAM) for 15 min on ice. Beads were wash once and eluted using loading dye (Biorad #161074).

### Mass spectrometry of kinase assays

After 1 hour incubation at 30⁰C, kinase reactions containing WT GST-SETD8 and GSK3α (Abcam ab42597) were reduced with 5mM DTT and alkylated with 20mM iodoacetamide. The samples were then digested with trypsin at a ratio of 1:50 (Promega #V5111) over night at room temperature. Prior to mass spectrometry analysis, samples were desalted using a 96-well plate filter (Orochem) packed with 1 mg of Oasis HLB C-18 resin (Waters). Briefly, the samples were resuspended in 100 µl of 0.1% TFA and loaded onto the HLB resin, which was previously equilibrated using 100 µl of the same buffer. After washing with 100 µl of 0.1% TFA, the samples were eluted with a buffer containing 70 µl of 60% acetonitrile and 0.1% TFA and then dried in a vacuum centrifuge. Samples were resuspended in 10 µl of 0.1% TFA and loaded onto a Dionex RSLC Ultimate 300 (Thermo Scientific), coupled online with an Orbitrap Fusion Lumos (Thermo Scientific). Chromatographic separation was performed with a two-column system, consisting of a C-18 trap cartridge (300 µm ID, 5 mm length) and a picofrit analytical column (75 µm ID, 25 cm length) packed in-house with reversed-phase Repro-Sil Pur C18-AQ 3 µm resin. Peptides were separated using a 90 min gradient from 4-30% buffer B (buffer A: 0.1% formic acid, buffer B: 80% acetonitrile + 0.1% formic acid) at a flow rate of 300 nl/min. The mass spectrometer was set to acquire spectra in a data-dependent acquisition (DDA) mode. Briefly, the full MS scan was set to 300-1200 m/z in the orbitrap with a resolution of 120,000 (at 200 m/z) and an AGC target of 5x10e5. MS/MS was performed in the ion trap using the top speed mode (2 secs), an AGC target of 1x10e4 and an HCD collision energy of 35.

Proteome raw files were searched using Proteome Discoverer software (v2.5, Thermo Scientific) using SEQUEST search engine and the SwissProt human database. The search for total proteome included variable modification of N-terminal acetylation, and fixed modification of carbamidomethyl cysteine. Trypsin was specified as the digestive enzyme with up to 2 missed cleavages allowed. Mass tolerance was set to 10 pm for precursor ions and 0.2 Da for product ions. Peptide and protein false discovery rate was set to 1%. Following the search, data was processed as previously described^62^. Briefly, proteins were log_2_ transformed, normalized by the average value of each sample and missing values were imputed using a normal distribution 2 standard deviations lower than the mean. Statistical regulation was assessed using heteroscedastic T-test (if p-value < 0.05). Data distribution was assumed to be normal but this was not formally tested.

### Generation of sgRNA vectors

The LentiCRISPRV2 (a gift from Feng Zhang, Addgene #52961), pSpCas9(BB)-2A-Puro (PX459) V2.0 (a gift from Feng Zhang, Addgene #62988), and LRG (a gift from Christopher Vakoc, Addgene #65656) vectors were used to generate gene-specific sgRNA vectors. For individual sgRNA cloning used in mESC engineering, PX459 was digested with BbsI (NEB #R3539S) and ligated with annealed sgRNAs targeting *Setd8* exon 4. For individual sgRNA cloning used in cell competition assays and phosphomutant rescue experiments, LRG and LentiCRISPRV2 were digested with BsmBI (NEB #R0739S) and ligated with annealed sgRNA oligos targeting *Setd8* exon 8. The sgRNA target sequences used included:

> *sgSetd8-1*:CTCAAGAAACGAACCGCCTA; *sgSetd8-2*: TAGGCGGTTCGTTTCTTGAG; *sgSetd8-e4*:GTAGAATCACATGATGGGGG; *sgNontargeting*: CATCATAAATGTACAACGGG

### Lentiviral preparation and infection

Viral vectors were prepped using ZymoPURE II™ Plasmid Maxiprep (Zymo Research D4202) and then transfected using PEI (Polysciences 49553-93-7) into HEK293T cells with packaging plasmids psPAX2 (Addgene #12260) and pMD2G (Addgene #12259). The viral supernatant was collected after 48 hours and 72 hours and passed through a 0.45µm filter. Lenti-X Concentrator (Takara #631231) was used to concentrate virus 50-100X. Monolayers were infected with lentivirus plus 1 μg/ml polybrene (Sigma-Aldrich #TR-1003-G) for 6h. Three days after the infection, monolayers were selected with Puromycin (10-12 µg/ml).

### Mouse models

All animals were maintained on a C57BL/6J background. C57BL/6J (JAX strain #000664) mice were obtained from The Jackson Laboratory. The Institutional Animal Care and Use Committee of the University of Pennsylvania (Animal Welfare Assurance Reference Number A3079-01, approved protocol no. 803415 granted to Dr. Lengner) approved all procedures involving mice and followed the guidelines set by the Guide for the Care and Use of Laboratory Animals of the National Research Council of the National Institutes of Health.

For phospho-mass spectrometry experiments, *Apc*^flox/flox58^::*Villin-CreERT* and *Gsk3α/β*^flox/flox23^::*Villin-CreERT* mice were injected intraperitoneally with 1mg tamoxifen dissolved in corn oil once per day for 3 days and collected 48h after the final injection. For azoxymethane (AOM)/ dextran sodium sulfate (DSS) experiments, *Setd8*^WT^ and *Setd8*^T138A^ mice were injected with 10mg/kg AOM (Sigma-Aldrich #A5486) intraperitoneally. 1 week later, 3X5 day cycles of 2- 2% DSS (MP Biomedicals #0216011050) in drinking water were provided to the mice, with two weeks between cycles. 3 weeks after the last cycle, mice were collected. The mice were closely monitored for signs of distress until the endpoint of the experiment.

### RT-PCR and qPCR

For gene expression analysis, RNA was isolated from tumoroids using TRIzol (Invitrogen# 15596026). 500 ng total RNA was used to synthesize cDNA using the High-Capacity cDNA Reverse Transcription Kit (Applied Biosystems #4368813). qPCR was performed using SYBR green (Applied Biosystems #4367659) on a QuantiStudio 7 Flex RealTime PCR machine (Applied Biosystems). Data was normalized by *Gapdh*. The following sets of primers were used to assess *Setd8* human cDNA mRNA expression levels:

> *SETD8*-human-cDNA-F1: 5’- CCGAACAAATGCTCTGGAATG -3’; *SETD8*-human-cDNA-R1: 5’-TTCCTGTAGATTCCGGCTAATG -3’

### Annexin V/PI staining

5 days post-transduction with LentiCRISPR-V2 containing *Setd8* sgRNAs, APK tumoroids were digested into single cells and stained with Dead Cell Apoptosis Kit with Annexin V Alexa Fluor 488 & PI (Invitrogen #V13241) according to the manufacturer’s instructions. An LSR Fortessa (BD Biosciences) was for flow cytometric analysis. The data was then quantified using Flowjo (BD Biosciences).

### EdU incorporation assays

5 days post-transduction with LentiCRISPR-V2 containing *Setd8* sgRNAs, 10 μM EdU was added to the monolayer culture media for 1.5 hours. Monolayers were digested into single cells and stained with eBioscience Fixable Viability Dye 780 (Invitrogen #65-0865-14). After, staining of EdU+ cells was performed with Click-iT Plus EdU Alexa Fluor 647 Flow Cytometry Assay Kit (Thermo Fisher #10634) according to the the manufacturer’s instructions. DNA was stained with Fxcycle Violet Stain (Thermo Scientific #F10347). An LSR Fortessa (BD Biosciences) was used to quantify the EdU incorporation and DNA content of live cells. The data was then analyzed using Flowjo (BD Biosciences).

### Clonal tumoroid formation assays

Tumoroids or monolayers were digested to single cells with Accutase and filtered with a 40µm filter. Viable cells were quantified using a hemocytometer and equal numbers of single cells were seeded in 24 well plates in complete medium supplemented with 10µM Y-27632. Brightfield images of each well were captured 5-7 days post-seeding with a Leica DM500 Widefield microscope and manually counted with Fiji.

### Bulk RNA-sequencing

4 days post-transduction with LentiCRISPR-V2 containing *Setd8* sgRNAs, RNA samples were extracted from WT and T140D SETD8 tumoroids using TRIzol (Invitrogen #15596026) according to the manufacturer’s instructions. RNA was then sent out for library preparation and next-generation sequencing by Novogene. Raw counts of transcripts were derived from fastq files mapped against Gencode M29 release with Salmon^63^ using standard settings. The resulting matrix was used as input for DESeq2 analysis^64^ to identify differentially expressed genes. Normalized counts outputted from DESeq2 was used as the input for GSEA^65^ for enrichment of gene signatures including fetal-like intestinal^41^ and YAP^66^ signatures. Differentially expressed genes (padj < 0.001 & log2FC > 1.0 or < -1.0) were inputted into EnrichR^67^ for pathway enrichment analysis.

### Cholesterol detection

4 days post-transduction with LentiCRISPR-V2 containing *Setd8* sgRNAs, SETD8 WT and T140D tumoroids were subjected to the Cholesterol/Cholesterol Ester-Glo™ assay (Promega #J3190) without esterase treatment according to the manufacturer’s instructions.

### ChIP-seq

4 days post-transduction with LentiCRISPR-V2 containing *Setd8* sgRNAs, WT and T140D SETD8 monolayers were washed with PBS then fixed with 1% formaldehyde for 10 minutes at room temperature. Formaldehyde was quenched with 125 mmol/L glycine. Purified nuclei isolated from fixed cells were sonicated with Covaris S220 sonicator with the following parameters: peak intensity, 140; duty factor, 5; cycles per burst, 200; time, 60 seconds on, 30 seconds off for 19 cycles. After centrifugation at 4 degrees with 13,500 rpm for 10 minutes, 5% of soluble chromatin was kept for input DNA and rest of chromatin was used to perform immunoprecipitation with 7.5μg of H4K20me1 antibody (Active Motif #39727) overnight at 4°C with rotation. Chromatin was then incubated with magnetic Protein G Dynabeads (Invitrogen #10003D) and then washed using low-salt, high-salt, LiCl buffer, and TE buffer. Bound chromatin was then eluted, reverse-cross-linked, and incubated with RNase A and Proteinase K. DNA samples were purified using the MinElute PCR purification kit (Qiagen #28004) and used for preparation of ChIP-seq libraries.

ChIP-seq libraries were prepped using the NEBNext Ultra II DNA Library Prep Kit (New England Biolabs E7645S) with NEBNext Multiplex Oligos Index Primers Sets for multiplexing. Libraries were sent to Novogene for sequencing. Sequencing reads were aligned to the mouse genome (mm10) using Bowtie2 with default settings^68^. Resulting SAM files were filtered for uniquely mapped reads and converted to BAM files, which were then sorted and marked for duplicate reads using SAMtools^69^. Technical replicates were then merged and peak calling was performed on merged BAM files using Epic2^70^ with default settings. Peak annotation was performed with ChIPseeker^71^ using UCSC mm10 known gene coordinates. Bigwig files were generated with deepTools^72^ bamcoverage with RPGC normalization and binSize of 50. Bigwig files were then normalized by input samples using bigwigCompare. Resulting bigwig files were subjected to K-means clustering analysis using deepTools computeMatrix scale-regions function utilizing BED files derived from genes expressed in APK tumoroids derived from bulk RNA-seq.

### TCGA survival

Colorectal cancer TCGA patients were scored according to their match to the expression pattern of the 50 genes most downregulated in SETD8-T140D vs. SETD8-T140A CAPK organoids using the singscore R package. A stronger signature match score indicates greater similarity to the SETD8-T140D expression state compared to the SETD8-T140A expression state. CRC TCGA Patients were then stratified into those with above median vs. below median SETD8-T140D signature scores and differences in overall survival between the two groups were evaluated using the survminer R package. Kaplan-Meier plots were generated and significance was tested via log-rank tests using the survdiff function.

### Mouse embryonic stem cell and genotyping SETD8 T138A

mESCs were nucleofected with 10μg PX459 containing *sgSetd8-e4* sgRNAs and 0.5nmol of single-stranded oligodeoxynucleotides (ssODNs) using Mouse ES Cell Nucleofector® Kit (Lonzo VPH-1001) according to the manufacturer’s instructions. mESCs were subsequently selected with 2μg/mL puromycin and subcloned.

The following ssODN sequences were used:

> *Setd8-T138A*:5’- GTTATACGAAGCGCTGTGAAGTCAGATGAACAGAAGAGCAAAGACACCAGGAGAGGTCCCCTGGCGCCTTTTCCAAACCAAAAATCCGAAGCTGCAGAACCTCCAAAAGCTCCACCTCCAAGTTGTGATTCTACCAATGTAGCAGTCGCTAAGCAAGCCCTGAAAAAGTCCCTCAAGGGCAAACAGGCCCCTCGGAAAAA-3’ *Setd8-T138D*: 5’- TTATACGAAGCGCTGTGAAGTCAGATGAACAGAAGAGCAAAGACACCAGGAGAGGTCCCCTGGCGCCTTTTCCAAACCAAAAATCCGAAGCTGCAGAACCTCCAAAAGATCCACCTCCAAGTTGTGATTCTACCAATGTAGCAGTCGCTAAGCAAGCCCTGAAAAAGTCCCTCAAGGGCAAACAGGCCCCTCGGAAAAAG-3’

For genotyping of T138A mESCs and mice, genomic DNA was amplified using Hot-Start Taq Blue (Thomas Scientific C775Y41) using the following primers:

> *Setd8-T138A-F*: 5’- GAAAAGGAACACCGGGAACGTTAT-3’ and *Setd8-T138A-R*: 5’- CCCAGGAACCAAGTGGTGTTAGG-3’.

Afterwards, the PCR product was digested overnight at 37⁰C with PstI (NEB R0140S) and the resulting digestion products were resolved on an agarose gel.

### MTT assay

WT or T138A colonic organoids were digested into single cells using Accutase and equal numbers of single cells were seeded in 24 well plates in complete medium supplemented with 10µM Y- 27632. 5 days after seeding, MTT reagents (Roche #11465007001) were added to each well according to manufacturer’s instructions, transferred to 96 well plate and read on SpectraMax 190 (Molecular Devices).

### Immunofluorescence

After dissection and washing with PBS, Swiss rolls of intestines or tumors were fixed in 4% paraformaldehyde (Electron Microscopy Sciences #15710), paraffin-embedded and sectioned. Hematoxylin & eosin staining was conducted by Morphology Core of the UPenn NIDDK P30 Center for Molecular Studies in Digestive and Liver Diseases. For immunofluorescence staining, antigen retrieval was performed in a pressure cooker using 0.01 M Tris-EDTA (pH 9.0) buffer. The sections were blocked with 10% goat serum and 1% BSA in TBS for 1 hour at room temperature and incubated with primary antibodies in 1% BSA in TBS overnight at 4°C. The next day, the sections were incubated with Cy2 and Cy3-conjugated fluorescent secondary antibodies (Jackson Laboratory, 1:500) and afterwards, DAPI in mounting media was added (Invitrogen #P36930). The following antibodies were used: anti-Ki67 (Abcam #ab15580, 1:200), and anti-E-cadherin (R&D Systems #AF748).

### Software and statistical analysis

R and PRISM software were used for statistical analysis, data visualization, and data processing. (http://www.graphpad.com). The R environment for graphics (https://www.r-project.org) was used in this study for the bioinformatics analysis of RNA-seq and ChIP-seq datasets. The R packages used for all analysis described in this manuscript were from the Bioconductor and CRAN, including enhancedVolcano^73^. enrichR^67^ was used for gene set enrichment analysis.

### Sequencing Data

Sequencing data for bulk RNA sequencing and ChIP-Sequencing can be found in the Gene Expression Omnibus (GEO) with accession numbers GSE270360 and GSE270361.

## Results

To identify GSK3 phosphorylation events that are regulated by APC, we conducted phospho-mass spectrometric profiling of murine intestinal epithelium after acute deletion of either *Apc* or *Gsk3α/β*^22,23^. Specifically, we crossed a *Villin-CreER* allele into *Apc*^flox/flox^ and *Gsk3α/β*^flox/flox^ strains to induce conditional knockout of these genes in the intestinal epithelium. We administered 3 daily doses of tamoxifen to these mice along with a control cohort (*Villin-CreER* only). 3 days after the final tamoxifen dose, intestinal crypts were collected and subjected to phosphoproteomic profiling. Successful recombination at each locus was verified by PCR (Fig S1A).

Considering all samples, we identified 8556 phosphosites belonging to 2734 unique proteins, the majority of which were serine and threonine residues (Fig 1A). Among these phosphosites, 1613 (18.8%) conformed to the GSK3 consensus motif (S/TXXXS/T) (Fig 1B). The proportion of phosphosites significantly decreased upon GSK3 loss (*p* < 0.05 & log_2_FC < -1) that conformed to the GSK3 consensus motif was significantly higher compared to all captured sites and those depleted upon *Apc* loss (Fig 1B and 1C). 205 and 147 phosphosites were significantly reduced upon *Gsk3α/β* and *Apc* loss, respectively, when compared to control (*p* < 0.05 & log_2_FC < -1) (Figs 1D and 1E). Differentially phosphorylated proteins from both genotypes were enriched for similar pathways, including Cadherin binding, RNA binding, estrogen response and TGF-β signaling (Figs 1F, G and S1B, C). 38 phosphosites were decreased upon both *Gsk3α/β* and *Apc* loss (Fig 1H), 7 of which conform to the GSK3 consensus motif (Fig S1D). These data indicate our experiment faithfully captured phosphoproteomic changes upon GSK3 loss and that APC and GSK3 are required for similar phosphorylation events in the intestinal epithelium.

**Figure 1.**
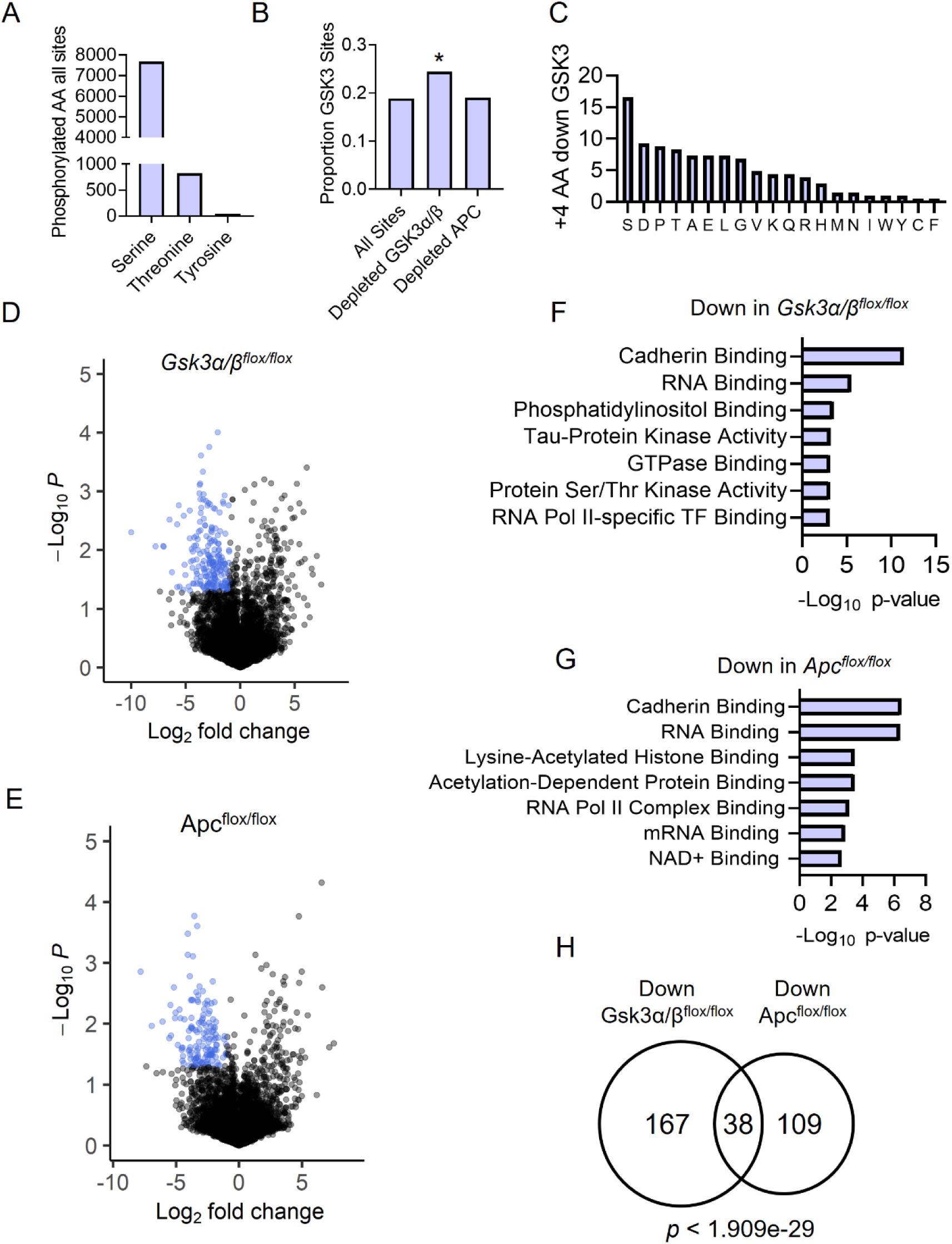
Phospho-mass spectrometry identifies candidate GSK3 phosphorylation events regulated by APC. **A.** Distribution of phosphorylated amino acids identified in phospho-mass spectrometry experiment. **B.** Proportion of phosphorylated residues residing within a GSK3 consensus recognition motif (S/TXXXS/T) within all phosphoproteins or phosphoproteins significantly reduced in *Gsk3α/β*^flox/flox^ or *Apc*^flox/flox^ intestines after *Villin-CreER*- mediated deletion, relative to control (*p* < 0.05 & log_2_FC < - 1). Statistical significance was determined by Chi-squared test and two-tailed heteroscedastic t-test. * *p* < 0.05. **C.** Percentage of amino acids +4 downstream of phosphosites reduced in *Gsk3α/β*^flox/flox^ intestines. **D, E.** Volcano plot of phosphoproteins in *Gsk3α/β*^flox/flox^ **(D)** or *Apc*^flox/flox^ intestines **(E)** relative to control. **F, G.** EnrichR pathway enrichment analysis of proteins reduced in *Gsk3α/β*^flox/flox^ **(F)** or *Apc*^flox/flox^ intestines **(G)** relative to control. **H.** Overlap of significantly reduced phosphoproteins (*p* < 0.05 & log_2_FC < -1). Statistical significance was assessed by a Hypergeometric test. (n = 2).

Among the phosphosites reduced after loss of both APC and GSK3 was Thr138 of the lysine methyltransferase SETD8 (Fig 2A). Although prior published phosphoproteomic studies identified phospho-Thr138 and its human homolog phospho-Thr140 in multiple cell types, the kinase mediating this event and its functional significance are not known (Fig 2B). This residue/recognition motif is highly conserved among amniotes, suggesting an evolutionarily conserved function (Fig 2C). Importantly, total SETD8 protein levels were not altered after loss of GSK3 or APC, indicating the loss of phospho-Thr138 is not a byproduct of total SETD8 reduction (Fig 2D). Therefore, we conclude that phosphorylation of SETD8 Thr138 in the intestinal epithelium is dependent on GSK3 and APC.

**Figure 2.**
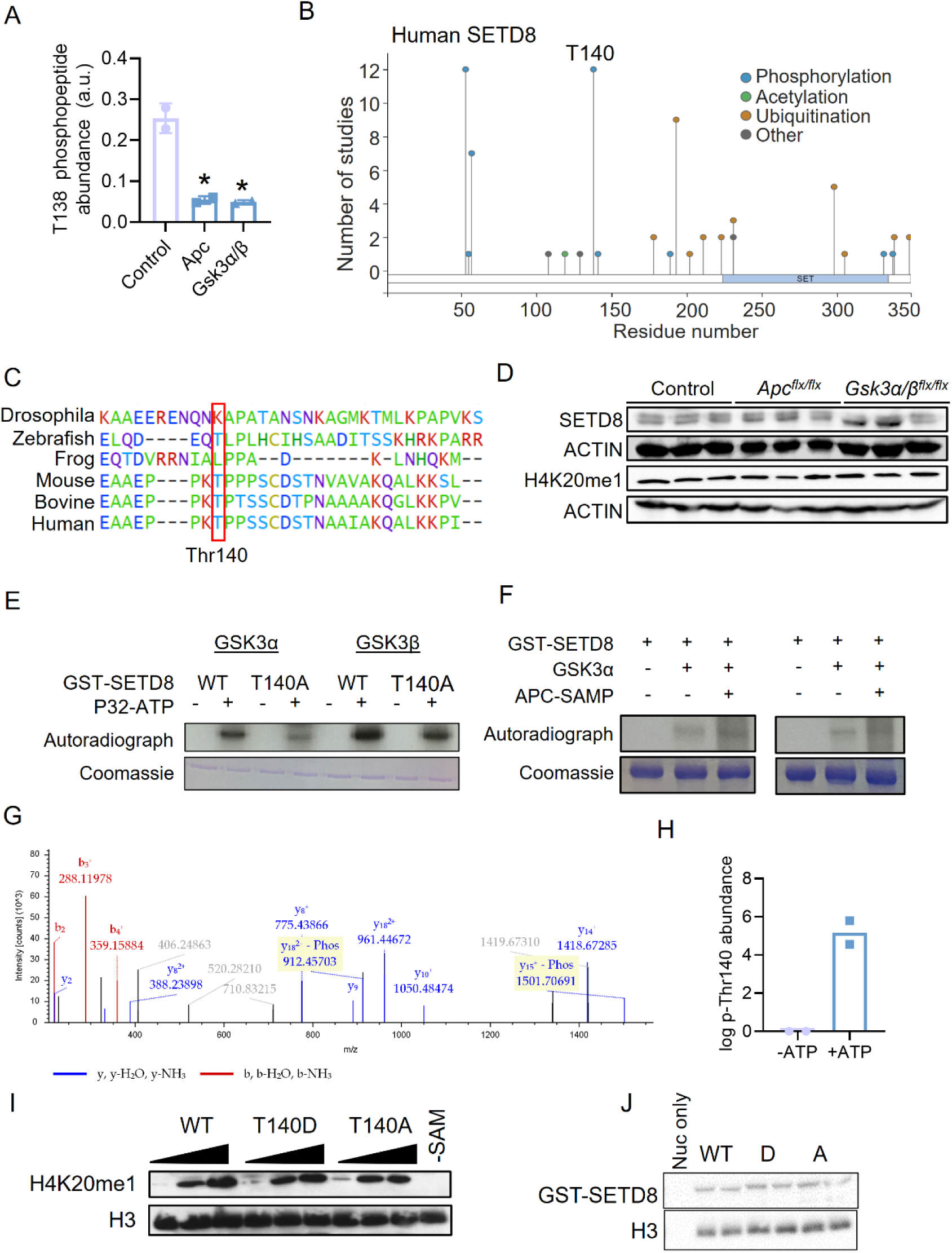
SETD8 Threonine 138 phosphorylation is reduced upon loss of APC or GSK3. **A.** Abundance of phospho-Thr138 SETD8 in control, *Gsk3α/β*^flox/flox^ or *Apc*^flox/flox^ intestines. Statistical significance was assessed by a two-tailed heteroscedastic t-test. (n = 2). * p < 0.05. **B.** The number of studies that have identified human SETD8 post-translation modifications from Phosphositeplus database. Threonine 140 is highlighted. **C.** Conservation of T140/T138 and adjacent amino acids between species. **D.** Western blot analysis of total SETD8 and H4K20me1 in control, *Gsk3α/β*^flox/flox^ and *Apc*^flox/flox^ intestines. β-ACTIN is used as a loading control. **E, F** Autoradiograph (top) and Coomassie stain (bottom) of recombinant radioactive kinase assays using GSK3α or GSK3β incubated with WT or T140A-SETD8 **(E)** and with or without recombinant APC SAMP region **(F)**. **G.** Annotated MS/MS spectrum of the peptide SEAAEPPKTPPSSCDSTNAAIAK modified with a phosphorylation on the T138 residue. Annotated spectrum indicates correct localization of the modification. Matching fragments from the N-terminus are highlighted in red (b series), while matching fragments from the C-terminus are highlighted in blue (y series). The subscript number indicates the fragment number, e.g. y2 is the fragment corresponding to the last two amino acids of the C-terminus. The superscript number corresponds to the charge state of the fragment ion. **H.** Abundance of the T140-phosphorylated SETD8 in kinase reactions with and without ATP. **I.** H3.1 mononucleosome methylation assay of wildtype or phosphomutant/mimetic T140 SETD8. H3 is included as a loading control. Reaction without S-adenosyl-methionine (- SAM) is included as a negative control. **J.** GST pull-down nucleosome binding assays of WT or phosphomutant/mimetic T140 SETD8 incubated with H3.1 mononucleosome. H3 is included as a loading control. Nucleosome only (nuc only) is included as a negative control.

To verify phosphorylation of Thr140 by GSK3/APC, we employed radioactive kinase assays. Human recombinant GSK3α and GSK3β both phosphorylated human SETD8 *in vitro* (Fig 2E). SETD8 T140A mutation reduced SETD8 phosphorylation by GSK3α and, to a lesser degree, GSK3β, suggesting that phosphorylation was specific for T140 (Fig 2E). Moreover, addition of an APC fragment containing SAMP repeats^5^ enhanced GSK3α phosphorylation of SETD8 (Fig 2F). Finally, mass spectrometry confirmed that GSK3α phosphorylated Thr140 *in vitro* (Fig 2G and 2H). We conclude that SETD8 Thr138/Thr140 is a bona fide substrate of GSK3 and that this phosphorylation event is potentiated by the presence of APC *in vitro* and *in vivo*.

SETD8 is the only lysine methyltransferase known to catalyze H4K20 monomethylation (H4K20me1), a histone modification important in myriad cellular functions, including cell cycle progression, gene expression and DNA damage repair^12^. We investigated whether Thr140 phosphorylation affected basic functionality of SETD8. We first found that loss of APC or GSK3 in the intestinal epithelium did not alter global levels of H4K20me1 (Fig 2D). We next generated recombinant SETD8 with phosphomutant (T140A) or phosphomimetic (T140D) mutations at T140. We found that neither alteration affected SETD8 enzymatic activity or nucleosome binding (Figs 2I and 2J). Furthermore, levels of total SETD8 and H4K20me1 were similar after expression of phosphomutant and phosphomimetic SETD8 T140 constructs in 293T cells (Fig S1E). Thus, Thr138/Thr140 phosphorylation does not affect SETD8 stability, methyltransferase function or nucleosome binding.

Given the importance of APC loss in CRC, we examined the contribution of SETD8 Thr140 phosphorylation to CRC phenotypes. To this end, we leveraged a genetically engineered murine colon tumor organoid (tumoroid) model bearing oncogenic mutations in *Apc*, *Kras*, and *Trp53*^24^ (*APK*). We first assessed the importance of total SETD8 in these tumoroids utilizing sgRNA competition assays. Consistent with an essential function in many types of cancers, cells transduced with sgRNAs targeting *Setd8* were quickly outcompeted by untransduced cells, in contrast to cells receiving control sgRNAs (Fig S2A). Moreover, we found that *Setd8* sgRNAs led to decreased tumoroid formation capacity and an induction of apoptosis without significantly altering proliferation (Figs S2B-G). Thus, dependency on SETD8 in this cellular context appears rooted in maintenance of self-renewal and survival pathways. Next, we leveraged sgRNAs targeting murine *Setd8* but not human *SETD8*, enabling the expression of human SETD8 mutants in the absence of endogenous murine SETD8 in *APK* tumoroids (Fig 3A). We derived stable *APK* lines expressing human WT, phosphomimetic T140D or phosphomutant T140A *SETD8* and then delivered sgRNAs targeting endogenous murine *Setd8* (Figs 3B and S3A-S3C). We observed that while WT and phosphomutant T140A SETD8 rescued self-renewal and viability phenotypes, phosphomimetic T140D did not (Figs 3C-3F). Additionally, phosphomimetic T140D did not alter *APK* tumoroid proliferation (Fig S3D). Therefore, the loss of Thr138/140 SETD8 phosphorylation, as observed downstream of APC inactivation, is critical for maintaining the self-renewal and viability of *APK* tumoroids.

**Figure 3.**
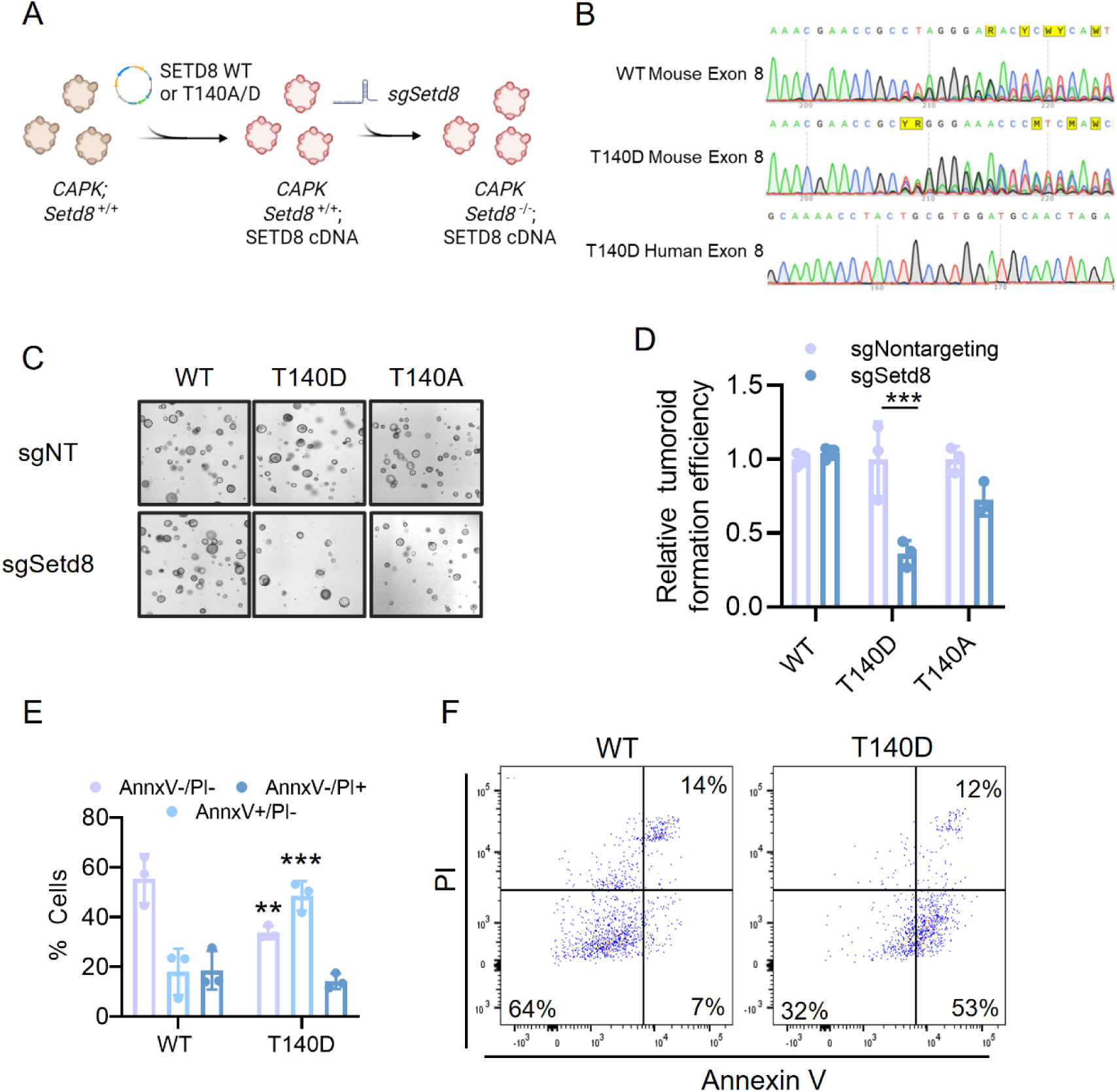
Loss of T140 SETD8 phosphorylation is critical in APC- mutant CRC. **A.** Schematic outline of SETD8 rescue experiments. Murine APK tumoroids were engineered to express human *SETD8* cDNAs with or without phosphomutant/mimetic (T140A/T140D) mutations at T140. sgRNAs targeting endogenous murine *Setd8* were then delivered into the resulting populations. **B.** Sanger sequencing tracks of sgRNA target region within mouse *Setd8* and homologous region of human *SETD8* after transduction with *Setd8* sgRNAs. **C, D.** Representative images **(C)** and quantification **(D)** of tumoroid formation assays of APK tumoroids expressing WT, T140D or T140A SETD8 and transduced with the indicated sgRNAs. NT: non-targeting. Statistical significance was assessed using an unpaired Student’s t-test. (n = 3). **E, F** Quantification **(E)** and representative flow plots **(F)** of Annexin-V and propidium iodide (PI) staining of APK tumoroids expressing WT or T140D SETD8 and transduced with sgRNAs targeting endogenous *Setd8*. Statistical significance was assessed using a two-way ANOVA with Holm-Šídák multiple comparison. (n = 3). For all panels: ** *p* < 0.01, *** *p* < 0.001.

SETD8 and H4K20me1 influence gene expression, with evidence supporting activating^25–28^, repressive^29–31^ and mixed^32–34^ roles in transcriptional regulation. We interrogated the effects of SETD8 Thr138/140 phosphorylation on the transcriptome of *APK* tumoroids using RNA-sequencing (Fig 4A)^12^. We observed mixed effects of Thr140 phosphorylation on gene expression; ∼700 and ∼500 genes were down- and up-regulated in SETD8 T140D (phosphomimetic) *APK* tumoroids, respectively (*p*_adj_ < 0.001 & log_2_FC > or < 1.0) (Fig 4B). Interestingly, genes significantly downregulated in T140D tumoroids were strongly enriched with gene sets relating to cholesterol biosynthesis and metabolism (Figs 4C-D and S4A). Maintaining cholesterol biosynthesis is critical for numerous cancer types^35,36^, including CRC^37–40^. Indeed, *Apc* loss in mice is directly linked to enhanced cholesterol biosynthesis^39^. Consistent with the transcriptional silencing of cholesterol biosynthetic enzymes, free intracellular cholesterol was significantly reduced in T140D tumoroids (Fig 4E). In addition, T140D tumoroids suppressed a fetal intestinal stem cell (ISC) signature originally defined as genes enriched in murine fetal intestinal spheroids relative to adult intestinal organoids (Figs 4F and S4B)^41^. This fetal gene expression program is associated with worse prognosis in CRC patients and regenerative programs in the adult mouse intestine post-injury^42–46^. The transcriptional effector of Hippo signaling, YAP, is a central regulator of the fetal ISC gene expression program^42^. Consistently, YAP target genes were depleted from tumoroids after rescue with T140D SETD8 (Fig 4G and S4C). To more directly examine the role of SETD8 Thr140 phosphorylation in human cancer, we investigated whether the transcriptional programs regulated by phosphorylation were associated with CRC patient prognoses. Indeed, a gene expression signature derived from genes altered by SETD8 Thr140 phosphorylation predicted more favorable outcomes in CRC patients, consistent with the tumor suppressive effect of SETD8 Thr140 phosphorylation (Fig 4H). Together, these findings suggest that maintenance of cholesterol biosynthesis and fetal intestinal stem cell gene expression programs may underpin the dependency of CRC on loss of SETD8 T138/T140 phosphorylation.

**Figure 4.**
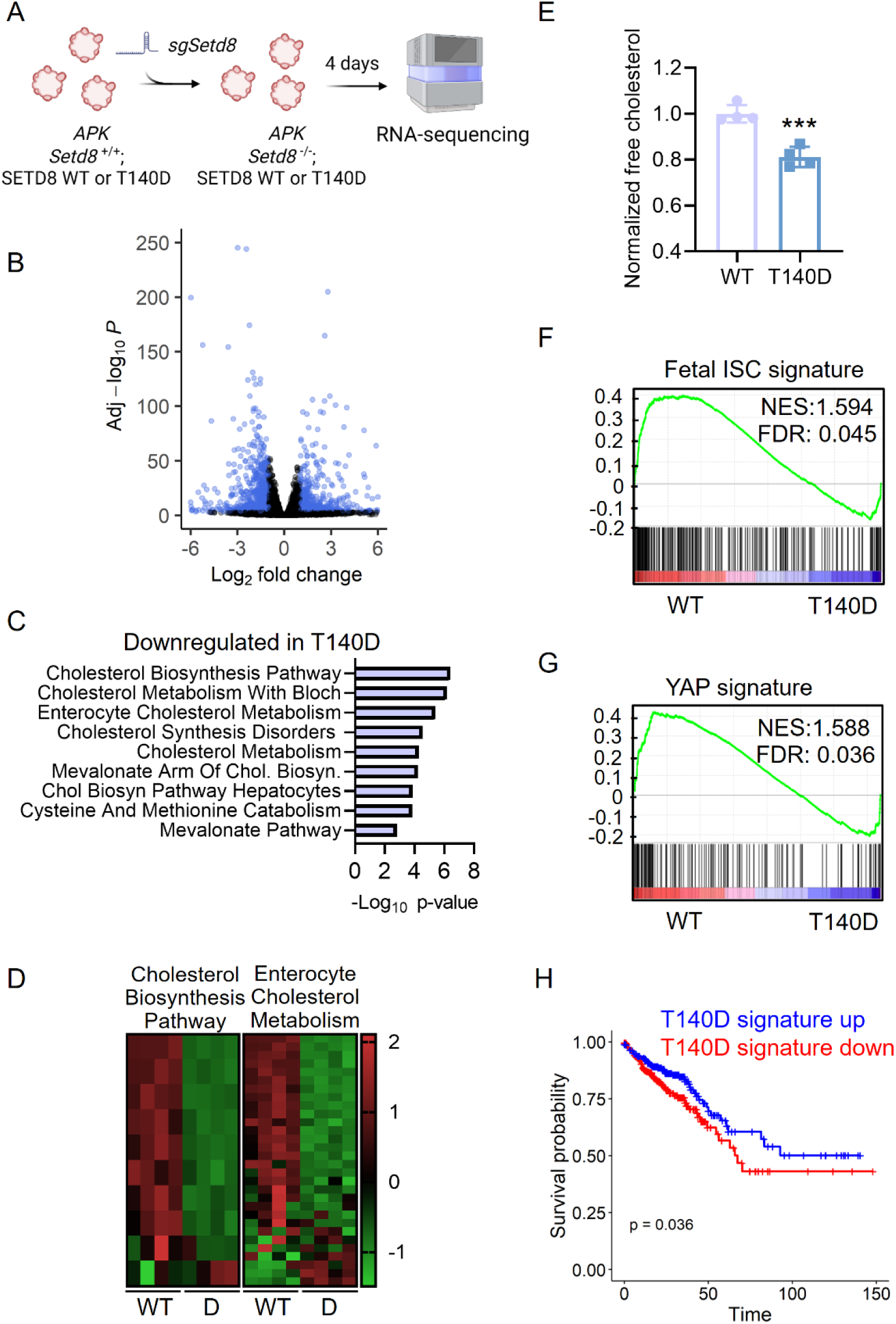
Phosphorylation of T140 SETD8 reprograms APC- mutant CRC gene expression. **A.** Overview of RNA-sequencing experiment assessing differences between WT and phosphomimetic T140D SETD8 function, **B.** Volcano plot of differential gene expression between WT and T140D SETD8 APK tumoroids. Statistical significance was determined by Wald’s test. Differentially expressed genes (*p*_adj_ < 0.001 & log_2_FC > 1.0 or < - 1.0) are highlighted. (n = 4). **C.** EnrichR results for genes downregulated in T140D tumoroids. **D.** Expression heatmap of z- scores of cholesterol-related gene sets in WT and T140D SETD8 APK tumoroids. **E.** Free cholesterol abundance in T140D and WT tumoroids normalized mean of WT controls. *** *p* < 0.001. **F, G.** Gene set enrichment plots of fetal-like **(F)** and YAP **(G)** signature genes in WT and T140D SETD8 APK tumoroids. **H.** Survival plots of COAD/READ patients stratified by SETD8 T140D signature.

We next investigated the role of SETD8 Thr140 phosphorylation on chromatin regulatory programs. Given previous difficulties in generating reliable SETD8 ChIP-seq^44^, as a proxy for SETD8 activity we mapped H4K20me1 in WT and T140D tumoroids collected under similar conditions as the previous RNA- sequencing experiment (Fig 4A). Consistent with previous studies^45^, H4K20me1 peaks were highly enriched within gene bodies (introns, exons, 5’ and 3’ UTR) and to a lesser extent promoter regions (Fig S4D). Also in line with previous findings, we found that H4K20me1 abundance correlated positively with gene expression globally (Figs S4E and S4F)^45^. We performed K-means clustering on gene body H4K20me1 signals and found several clusters with varying H4K20me1 patterns (Fig 5A). Notably, cluster 3 contained an overrepresentation of genes differentially expressed between WT and T140D organoids (Fig 5B). Interestingly, genes within this cluster that were downregulated in T140D organoids were enriched for cholesterol metabolism pathways (Fig 5C). This included enterocyte cholesterol genes, which were both downregulated and had reduced gene body H4K20me1 in T140D organoids (Figs 5D and 5E). Also enriched in this cluster were genes driving the fetal intestinal and related YAP pathway genes. These genes were downregulated in T140D organoids and, interestingly, had increased gene body H4K20me1 (Figs 5F-J and S4G). This suggests that Thr140 phosphorylation significantly affects H4K20me1 dynamics, leading to losses in oncogenic transcriptional programs such as cholesterol metabolism genes and YAP-driven fetal-like program.

**Figure 5.**
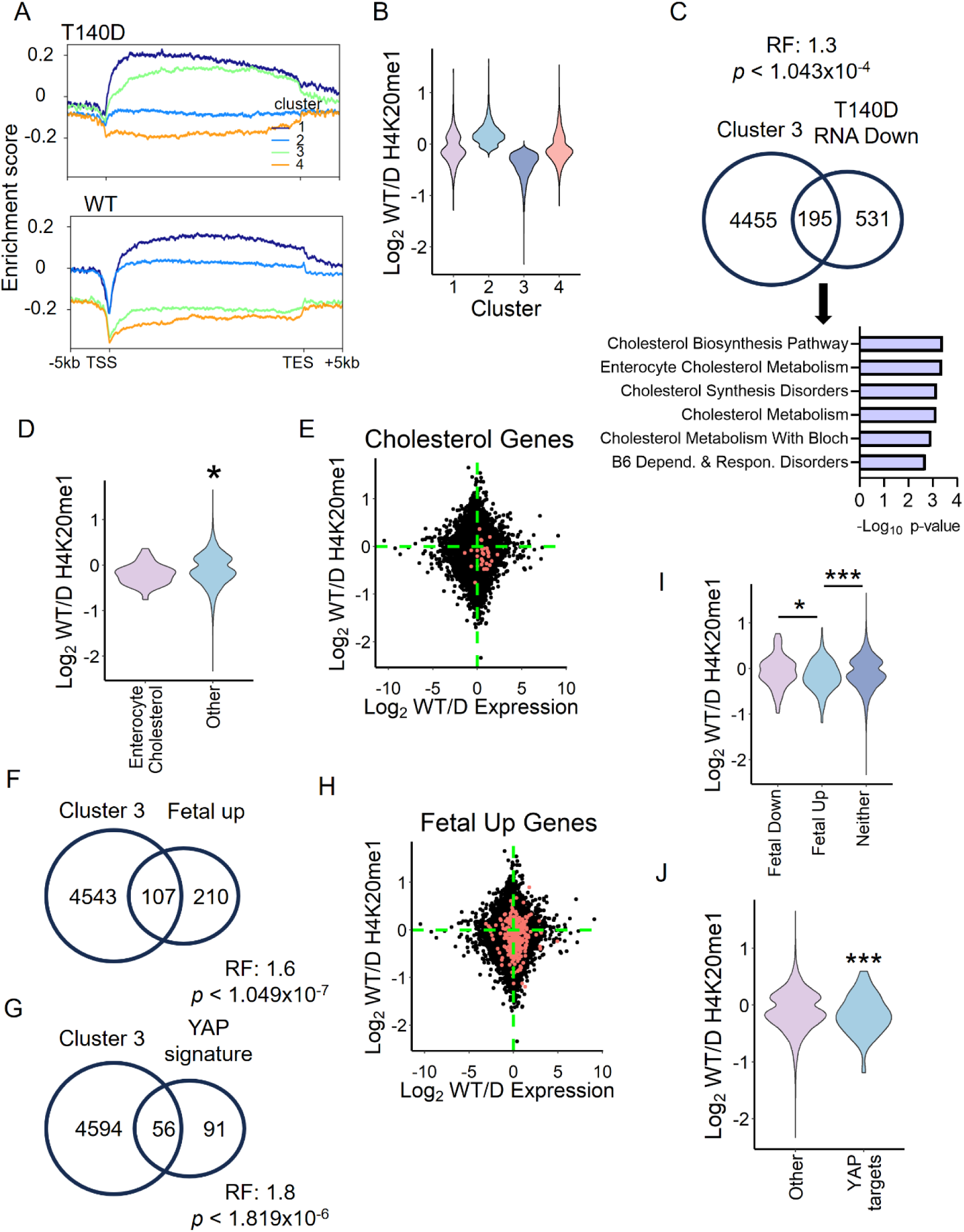
SETD8 T140 phosphorylation alters H4K20me1 localization. **A.** K-means clustering of H4K20me1 distribution within gene bodies in WT and T140D tumoroids. (n = 2). **B.** Log_2_FC of H4K20me1 between WT and T140D tumoroids across k-means clusters. **C.** Overlap between cluster 3 genes and genes significantly downregulated in T140D tumoroids (*p*_adj_ < 0.001 & log_2_FC < -1.0). Statistical significance was assessed by a Hypergeometric test. **D.** Log_2_FC of H4K20me1 between WT and T140D tumoroids of genes in enterocyte cholesterol geneset (wikipathway WP5333) compared to all other genes. **E.** Log_2_FC of H4K20me1 and gene expression between WT and T140D tumoroids, enterocyte cholesterol genes are highlighted in red. **F, G.** Overlap between cluster 3 genes and either genes upregulated in fetal intestines **(F)** or YAP signature genes **(G)**. Statistical significance was assessed by a Hypergeometric test. **H.** Log_2_FC of H4K20me1 and gene expression between WT and T140D tumoroids, fetal intestine genes are highlighted in red. **I, J.** Log_2_FC of H4K20me1 between WT and T140D tumoroids of genes upregulated or downregulated in fetal intestine **(I)** or YAP target genes **(J)** compared to all other genes. For all panels: * *p* < 0.05, *** *p* < 0.001. For panels 5D and 5J, statistical significance was determined by a Student’s t-test. For panel 5I, statistical significance was determined by a one-way ANOVA with Tukey’s multiple comparison. (n = 2).

Tumoroid experiments and epigenomic trends strongly suggest the importance of SETD8 T138/T140 phosphorylation in APC-mutant CRC. However, exogenous expression of SETD8 mutant constructs may not faithfully reflect endogenous SETD8 phosphorylation. To better model the consequences of SETD8 phosphorylation *in vivo*, we generated genetically engineered mice bearing phosphomutation at the endogenous *Setd8* locus. We designed sgRNAs and donor oligonucleotides to generate either phosphomutant (Thr->Ala) or phosphomimetic (Thr->Asp) mutations (Fig S5A). Interestingly, while we recovered 3/44 (6.8%) mESC clones with correctly targeted phosphomutant mutations, we were unable to recover any viable mESC clones with phosphomimetic mutations (Fig S5B). This trend approached statistical significance, suggesting that phosphomimetic T138D mutations may be incompatible with mESC viability, which would be in line with our findings in tumoroids that SETD8 phosphorylation suppresses stemness and induces apoptosis. Clones bearing T138A mutations were verified with Sanger sequencing and used to establish a SETD8 phosphomutant mouse colony (Figs S5C- E). *Setd8^T^*^138^*^A/T^*^138^*^A^* mice were viable and did not present any gross abnormalities during development and adulthood. Furthermore, SETD8 and H4K20me1 protein levels were unchanged in intestinal and colonic tissue of adult mice (Fig S5F). Additionally, we did not observe any changes in intestinal architecture or proliferation (Fig S5G). Colonic organoids derived from *Setd8^T^*^138^*^A/T^*^138^*^A^* mice exhibited similar organoid formation rates and proliferation compared to S*etd8*^WT/WT^ littermates (Fig S5H-S5J). These data indicate that loss of SETD8 T138 phosphorylation alone is insufficient to induce phenotypic changes in murine intestinal and colonic epithelium.

Given that regulation of SETD8 T138 phosphorylation is a downstream consequence of *Apc* inactivation, we reasoned that this phosphorylation event may be more relevant in an oncogenic context with additional genetic insults. To test this, we utilized the azoxymethane (AOM)/dextran sodium sulfate (DSS) protocol as a model for *de novo* CRC tumorigenesis. AOM is a mutagen whereas DSS disrupts the intestinal barrier, causing inflammation and generating colitis- associated cancers in the distal colon/rectum. These tumors do not carry frequent APC loss of function mutations^47^, and thus we reasoned that the Setd8 T138A phosphomutation may sensitize mice to AOM-DSS-initiated tumorigenesis. Indeed, S*etd8*^T138A/T138A^ mice were more susceptible to AOM/DSS tumorigenesis, bearing significantly more and larger tumors (Fig 6A- 6F). Therefore, loss of SETD8 Thr138 phosphorylation increases susceptibility to de novo CRC tumorigenesis, indicating that this post-translation modification is tumor- suppressive in CRC.

**Figure 6.**
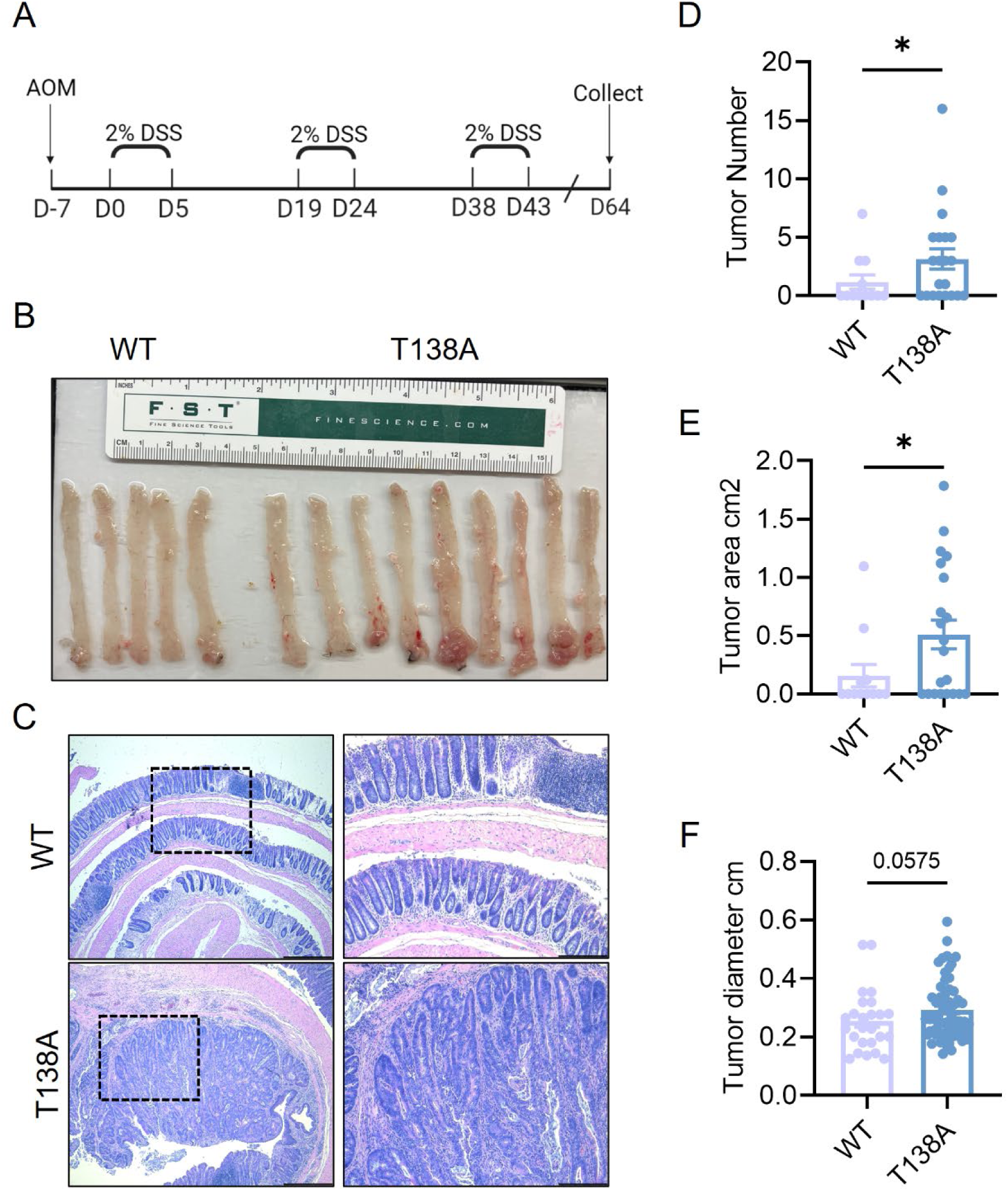
Setd8^T138A^ mice are more susceptible to azoxymethane (AOM)/ dextran sodium sulfate (DSS)-induced tumorigenesis. **A**. Timeline of AOM/DSS induction protocol. **B.** Representative images of colorectum from Setd8^WT^ and Setd8^T138A^ mice after AOM/DSS regimen. **C.** Representative hematoxylin and eosin staining of Setd8^WT^ and Setd8T138A mouse colorectum after AOM/DSS experiment. **D-F.** Quantification of tumor number **(D)**, tumor area **(E)** and tumor diameter **(F)** in Setd8^WT^ and Setd8^T138A^ mouse colorectum after AOM/DSS regimen. Statistical significance was determined using a Mann- Whitney U test. * p < 0.05. (n ≥ 12).

In total, we describe a novel APC/GSK3-dependent phosphorylation event in the adult intestinal epithelium whose loss is critical to maintain oncogenic gene expression programs in CRC.

## Discussion

Loss-of-function mutations in *APC* are the most frequent genetic aberration in CRC patients. Seminal work has defined the critical roles that *APC* plays in the destruction complex to coordinate the phosphorylation and degradation of β-CATENIN^3^. Nevertheless, the overrepresentation of *APC* loss-of-function mutations compared to gain-of-function mutations in *CTNNB1* (encoding β- CATENIN) suggest that the function of *APC* in colorectal epithelium goes beyond its role in regulating β-CATENIN^2^. Indeed, targeted studies have identified other GSK3 targets that are regulated by APC^5^.

Here, we systematically defined candidate GSK3 phosphorylation events mediated by APC *in vivo*. We identify SETD8 T138/T140 as a tumor suppressive phosphorylation event mediated by GSK3 and regulated by APC. Thus, in contrast to the prevailing dogma regarding the role of APC in CRC tumorigenesis, we find that APC loss dysregulates other important GSK3 targets beyond β-CATENIN. These data emphasize that the molecular changes that accompany APC loss are more complex than previously believed. Further identification of other GSK3 targets regulated by APC will enable a deeper understanding of APC loss in CRC.

We find that GSK3 directly phosphorylates SETD8 T138/T140 and that APC enhances this phosphorylation event. However, further work is needed to further characterize APC-driven GSK3 phosphorylation of SETD8. It is unknown whether AXIN and the other components of the destruction complex are also involved in SETD8 T138/T140 phosphorylation. Moreover, the upstream signaling pathways that dictate SETD8 T138/T140 phosphorylation in the absence of APC mutations are unclear. Past studies have found that some APC/GSK3 phosphorylation events require AXIN and are therefore responsive to extracellular WNT signals due to the recruitment of AXIN to LRP5/6^5^. Nevertheless, not every GSK3/APC-mediated phosphorylation event is AXIN-dependent/WNT-responsive, indicating that SETD8 T138/T140 phosphorylation could be regulated by other upstream signaling pathways^5^.

SETD8 is the only known enzyme that catalyzes monomethylation of H4K20 to H4K20me1. Additionally, SETD8 methylates non-histone substrates such as P53 and CD147^13,14^. SETD8 is overexpressed and essential in multiple cancer types, including CRC^13,19–21^. Nevertheless, many of these studies fail to define the underpinnings of cancer-specific changes in SETD8 regulation and function. This point is particularly salient given the common essentiality of SETD8 in human cell lines, which indicates that SETD8 is functionally required in mammalian cells regardless of oncogenic transformation. We find that phospho-T138/140 SETD8 is dramatically reduced upon loss of tumor suppressor *APC* and that this change in phosphorylation is critical in CRC tumorigenesis. We show that expression of T140D phosphomimetic SETD8 leads to drastic transcriptional changes in CRC tumoroids. Interestingly, these alterations included loss of a fetal-like gene expression program as well as cholesterol biosynthesis genes in T140D tumoroids. Activation of a fetal-like signature gene expression program has emerged as a critical event in CRC tumorigenesis^43–46^. This study provides insight into the molecular events underpinning the reactivation of fetal-like gene expression modules after *APC* loss. Future studies are needed to decipher the interplay between SETD8 with SOX9^44^ and YAP^42^, transcription factors known to drive the fetal gene expression program.

While global alterations in H4K20me1 methylation did not correlate with gene expression changes, loss of fetal and cholesterol biosynthesis gene expression was associated with gains in H4K20me1. These data are in alignment with other studies identifying a negative correlation between H4K20me1 and gene expression^29–31^. Future studies are needed to decipher why H4K20me1 can have mixed effects on gene transcription depending on the cellular and genomic context. Further, SETD8 could influence this gene expression program by methylation of non-histone substrates. Indeed, SETD8 was recently shown to drive YAP signaling by methylation of SNIP1^19^.

Biochemical and structural studies have yielded insight into SETD8 function^12^. However, structural studies often have focused on fragments of SETD8 containing only the C-terminal SET domain^48–51^. Notably, the C-terminal SET domain is sufficient for methylation of nucleosome substrates^51,52^. The poorly characterized N-terminal region of SETD8 is largely a disordered region. The only functions attributed to the N-terminal region of SETD8 are the regulation of SETD8 stability and protein-protein interactions with non-histone subtrates^53–56^. Our identification of T138/T140 as a functional phosphorylation target sheds light on the functionality of the N-terminal region of SETD8, but also highlights the need for further investigation of this region. Indeed, although we observed clear changes in H4K20me1 localization after expression of phosphomimetic SETD8 T140D, the direct consequences of T138/T140 phosphorylation on the SETD8 protein itself remain elusive, as this phosphorylation does not affect SETD8 catalytic activity, nucleosome binding, or protein stability. To this point, phosphorylation events are enriched in disordered regions of some histone methyltransferases/demethylases^57^, and this often initiates the formation of stable configurations that mediate protein-protein interactions^57^. Therefore, phosphorylation of SETD8 T138/T140 phosphorylation may regulate SETD8/H4K20me1 localization through affecting interaction with co-factors/transcription factors at specific gene loci.

## Supporting information

All figures

## Authors’ Disclosures

The authors have no financial interests related to this work to declare.

## Authors’ Contributions

**Z. Cramer:** Conceptualization, formal analysis, validation, investigation, visualization, methodology, writing–original draft, writing–review and editing. **K. Monaghan**: investigation, methodology. **R. Petroni**: investigation, methodology. **X. Wang**: investigation, methodology. **S. Adams-Tzivelekidi**s: Resources, project administration. **K. Durning**: investigation, methodology. **M. Kim**: visualization, methodology. **Y. Tian**: investigation, methodology. **N. Johnson**: investigation, methodology. **N. Leu**: investigation, methodology. **S. Sidoli**: investigation, methodology, visualization, formal analysis. **N. Li:** Conceptualization, formal analysis validation, investigation, methodology. **M. Blanco**: Conceptualization, supervision, funding acquisition, writing–original draft, writing–review and editing. **C. Lengner**: Conceptualization, supervision, funding acquisition, writing–original draft, writing–review and editing

## Acknowledgements

We thank Dr. Peter Klein for valuable discussions, both technical and conceptual, related to this work. We acknowledge the Penn Vet Comparative Pathology Core pathology core for analysis of *Setd8* phosphomutant mouse tissues, the Penn flow cytometry core, the Center for Molecular Studies in Digestive and liver Diseases (NIH P30-DK050306) and its Molecular Pathology and Imaging Core, and the Penn Vet Imaging Core (Gordon Ruthel). This work was supported by the following grants: NIH F31CA250267 (ZC), NIH/NCI R01 CA279317-01 (MAB), and NCI R01CA168654 (CJL).

## Notes

### Competing Interest Statement

The authors have declared no competing interest.

