## Supplementary material for "APC coordinates GSK3 phosphorylation of SETD8 to suppress colorectal cancer": All figures

Figure 1

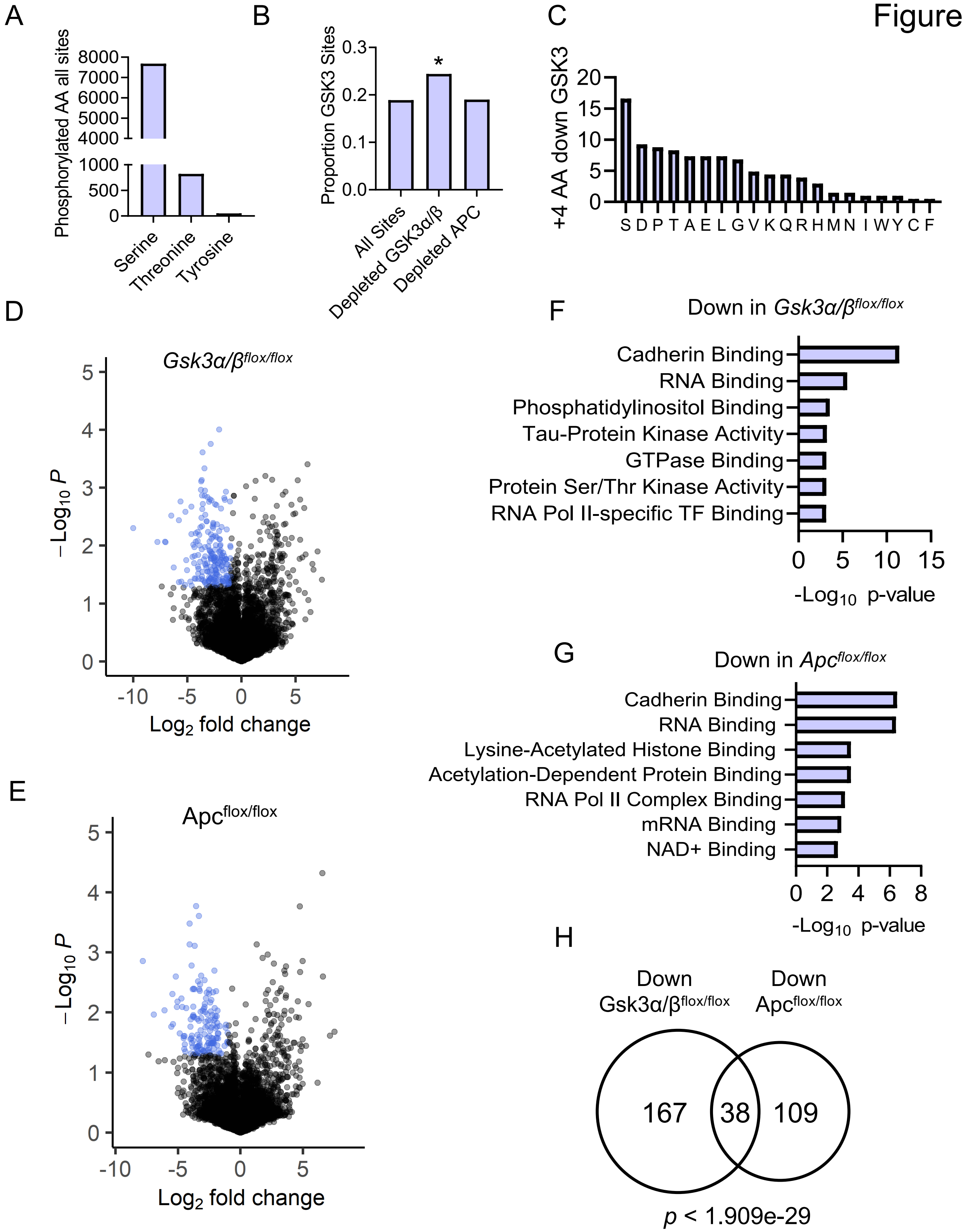

A
Phosphorylated AA all sites
8000
7000
6000
5000
4000
1000
500
0
Serine
Threonine
Tyrosine
B
Proportion GSK3 Sites
0.3
0.2
0.1
0.0
*
All Sites
Depleted GSK3α/β
Depleted APC
C
+4 AA down GSK3
20
15
10
5
0
S D P T A E L G V K Q R H M N I W Y C F
D
Gsk3α/β flox/flox
−Log10 P
Log2 fold change
E
Apc flox/flox
F
Down in Gsk3α/β flox/flox
Cadherin Binding
RNA Binding
Phosphatidylinositol Binding
Tau-Protein Kinase Activity
GTPase Binding
Protein Ser/Thr Kinase Activity
RNA Pol II-specific TF Binding
-Log10 p-value
G
Down in Apc flox/flox
Cadherin Binding
RNA Binding
Lysine-Acetylated Histone Binding
Acetylation-Dependent Protein Binding
RNA Pol II Complex Binding
mRNA Binding
NAD+ Binding
-Log10 p-value
H
Down
Gsk3α/β flox/flox
Down
Apc flox/flox
167
38
109
p < 1.909e-29

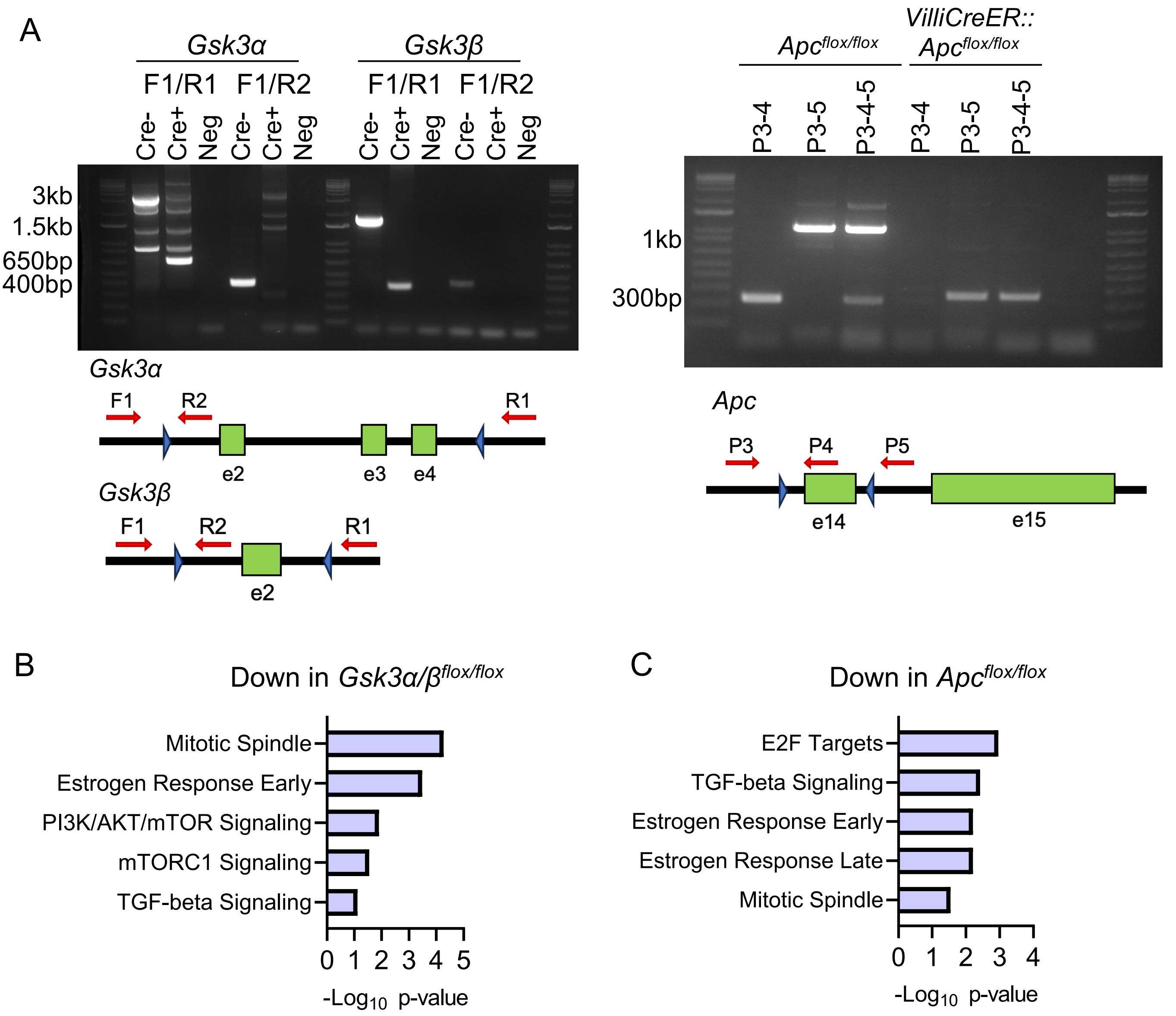


D

| Accession | Gene Name | Phosphosite |
|---|---|---|
| A0ELI5 | Edc3 | S161 |
| G3UWR2 | Gigyf2 | S19 |
| A0A0G2JFK5 | Setd8 | T138 |
| Q4VA98 | Zfp219 | S665 |
| O08715-2 | Akap1 | S64 |
| Q3TUQ7 | Prkaa1 | S477 |
| Q3TN42 | Utp15 | T515 |

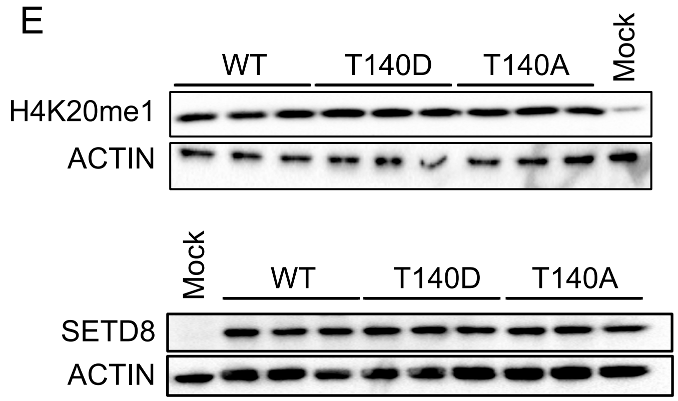


Supplemental Figure 1

Figure 2

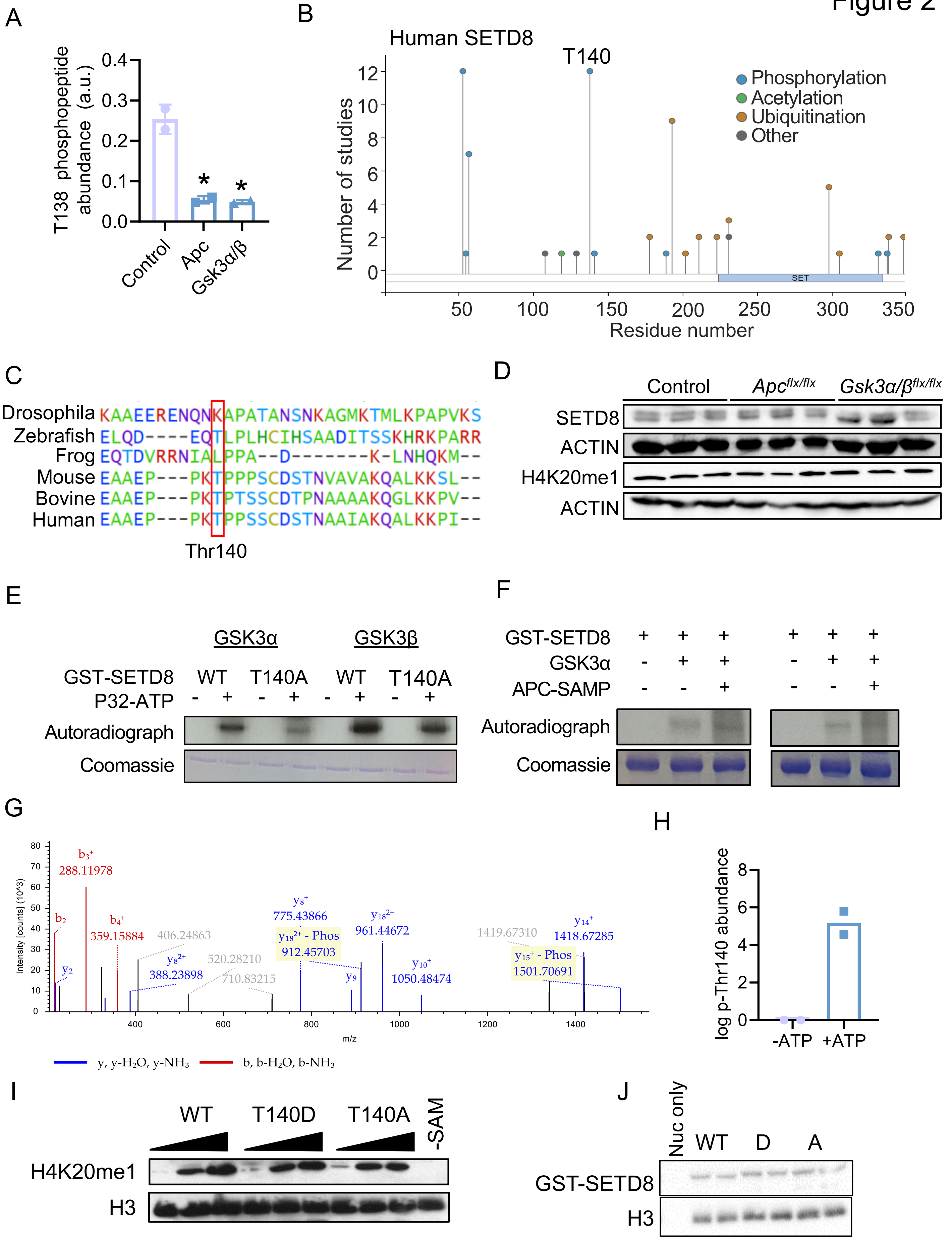
A
T138 phosphopeptide abundance (a.u.)
0.0
0.1
0.2
0.3
0.4
Control
Apc
Gsk3α/β
B
Human SETD8
T140
Phosphorylation
Acetylation
Ubiquitination
Other
Number of studies
SET
Residue number
C
Drosophila KAAEERENQNKAPATANSNKAGMKTMLKPAPVKS
Zebrafish ELQD----EQTLPLHCIHSAADITSSKHRKPARR
Frog EQTDVRRNIALPPA--D-------K-LNHQKM--
Mouse EAAEP---PKTPPPSCDSTNVAVAKQALKKSL--
Bovine EAAEP---PKTPTSSCDTPNAAAAKQGLKKPV--
Human EAAEP---PKTPPSSCDSTNAAIAKQALKKPI--
Thr140
D
Control
Apcflx/flx
Gsk3α/βflx/flx
SETD8
ACTIN
H4K20me1
ACTIN
E
GSK3α
GSK3β
GST-SETD8
WT
T140A
P32-ATP
Autoradiograph
Coomassie
F
GST-SETD8
GSK3α
APC-SAMP
Autoradiograph
Coomassie
G
Intensity [counts] (10^3)
m/z
y, y-H2O, y-NH3
b, b-H2O, b-NH3
H
log p-Thr140 abundance
-ATP
+ATP
I
WT
T140D
T140A
-SAM
H4K20me1
H3
J
Nuc only
WT
D
A
GST-SETD8
H3

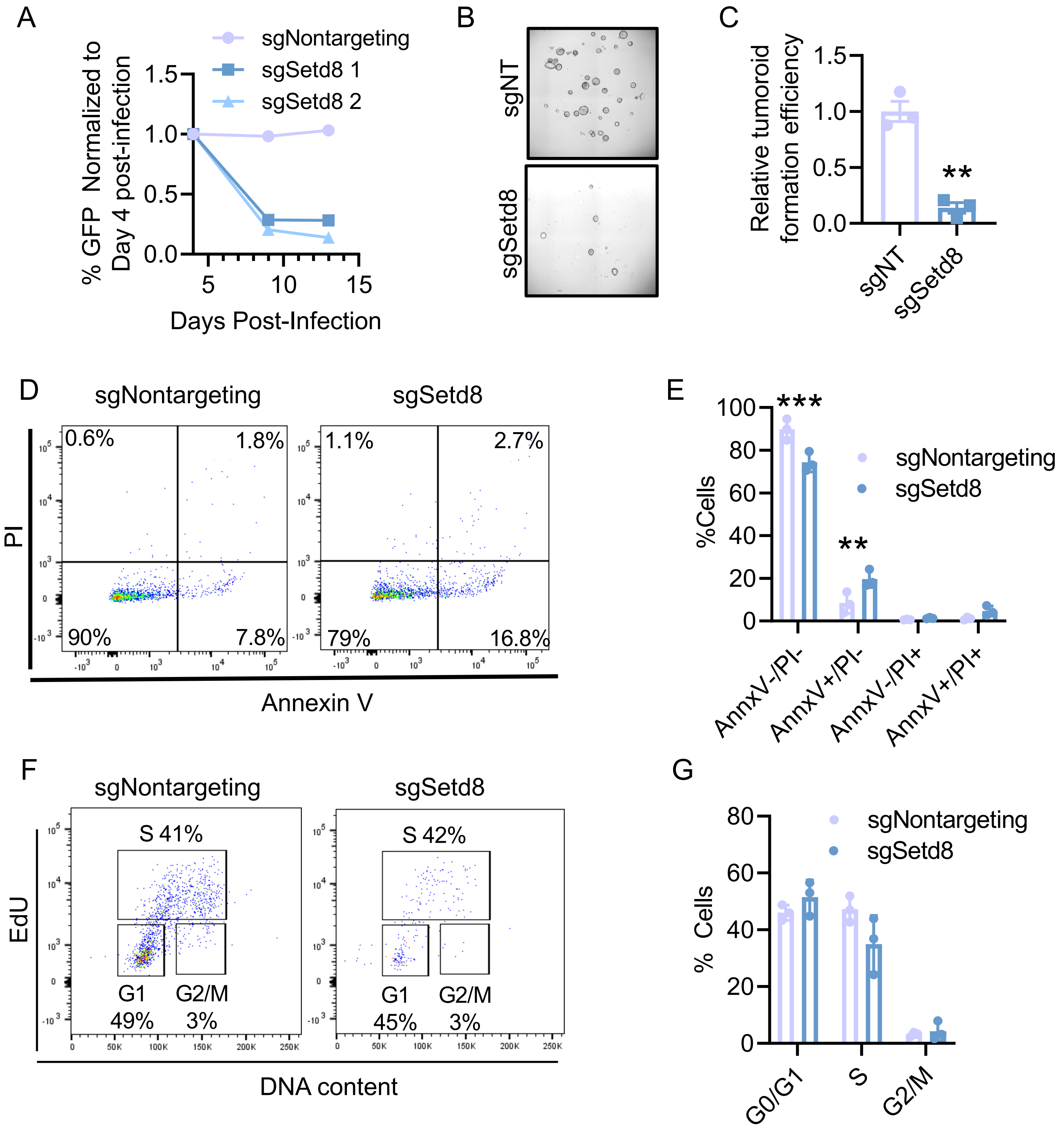


Supplemental Figure 2

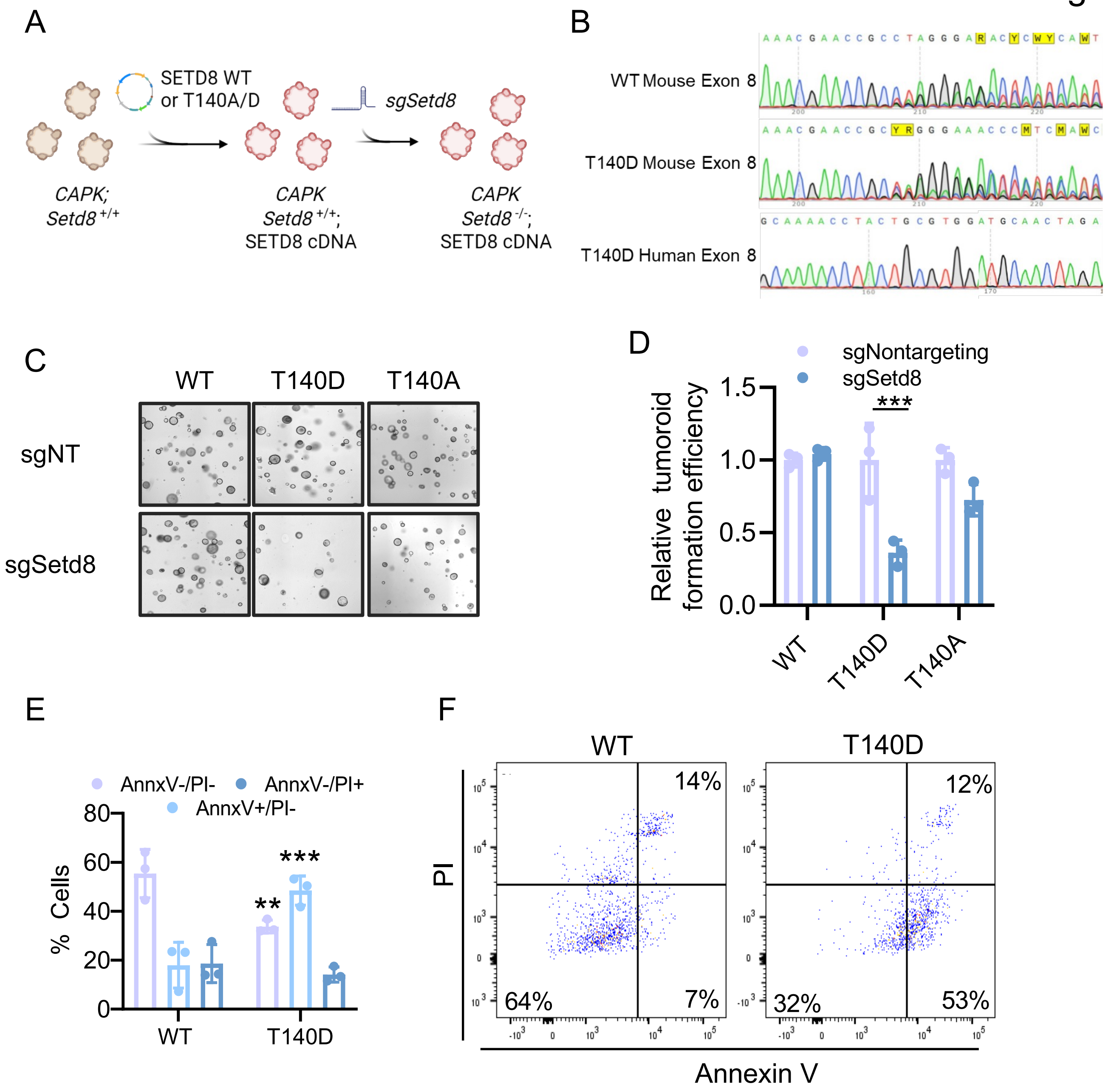
A
SETD8 WT
or T140A/D
sgSetd8
CAPK;
Setd8 +/+
CAPK
Setd8 +/+;
SETD8 cDNA
CAPK
Setd8 -/-;
SETD8 cDNA
B
WT Mouse Exon 8
T140D Mouse Exon 8
T140D Human Exon 8
C
WT
T140D
T140A
sgNT
sgSetd8
D
sgNontargeting
sgSetd8
Relative tumoroid formation efficiency
1.5
1.0
0.5
0.0
***
WT
T140D
T140A
E
AnnxV-/PI-
AnnxV-/PI+
AnnxV+/PI-
% Cells
80
60
40
20
0
**
***
WT
T140D
F
WT
T140D
PI
14%
64%
7%
12%
32%
53%
Annexin V

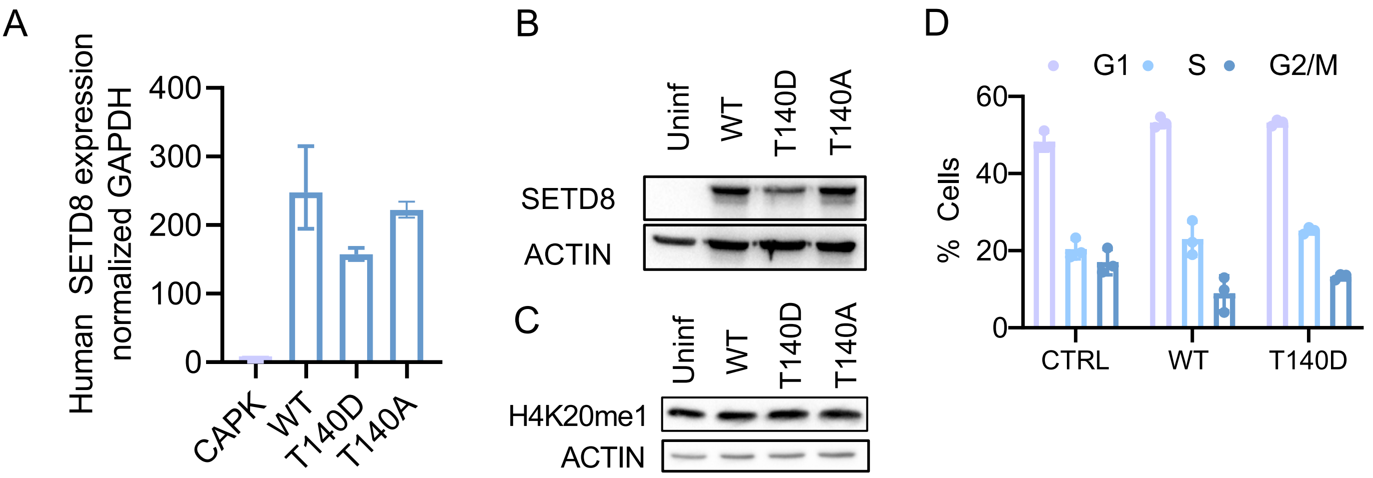


Supplemental Figure 3

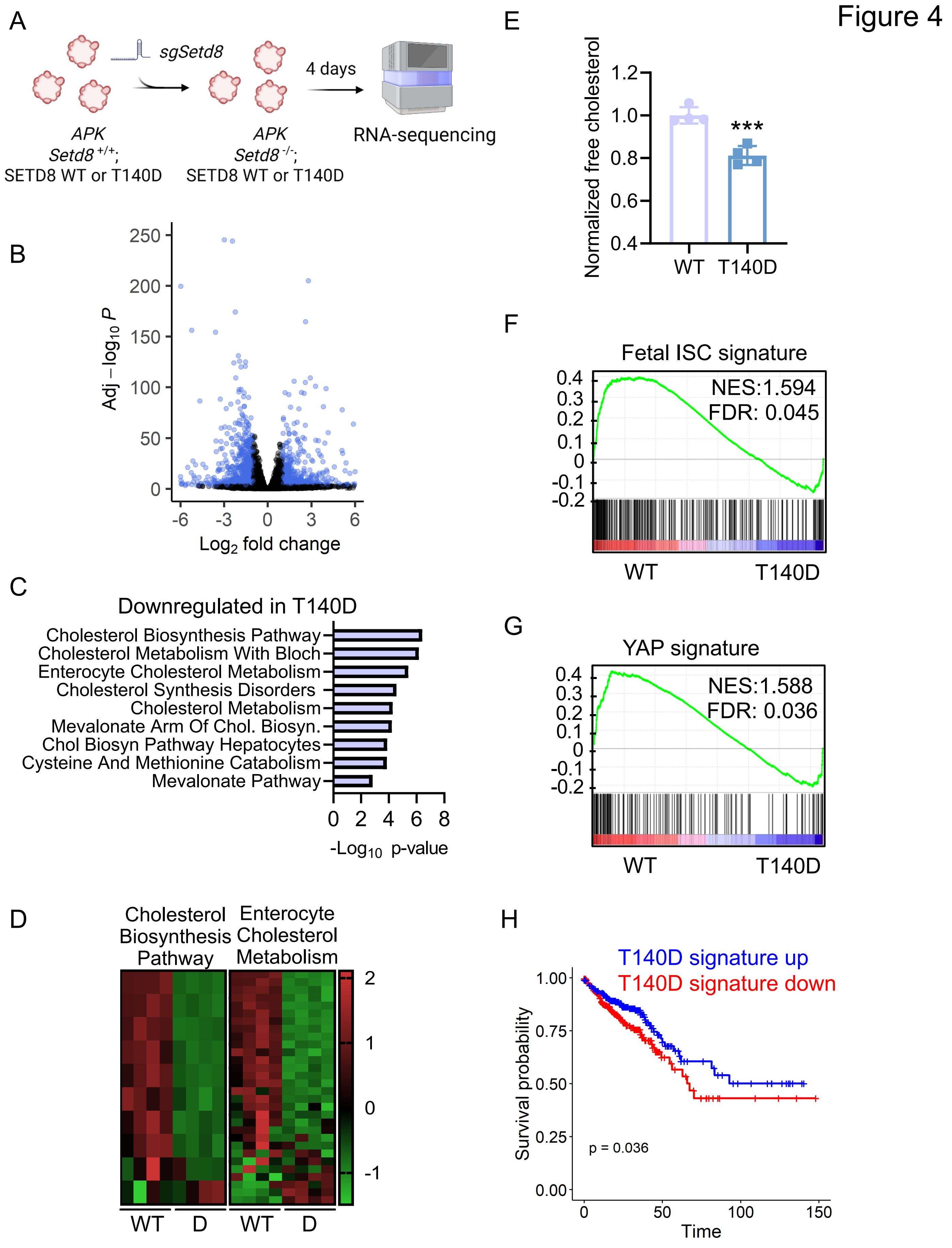
Figure 4
A
sgSetd8
4 days
APK
Setd8 +/+;
SETD8 WT or T140D
APK
Setd8 -/-;
SETD8 WT or T140D
RNA-sequencing
B
Adj −log10 P
Log2 fold change
C
Downregulated in T140D
Cholesterol Biosynthesis Pathway
Cholesterol Metabolism With Bloch
Enterocyte Cholesterol Metabolism
Cholesterol Synthesis Disorders
Cholesterol Metabolism
Mevalonate Arm Of Chol. Biosyn.
Chol Biosyn Pathway Hepatocytes
Cysteine And Methionine Catabolism
Mevalonate Pathway
-Log10 p-value
D
Cholesterol Biosynthesis Pathway
Enterocyte Cholesterol Metabolism
WT
D
WT
D
E
Normalized free cholesterol
***
WT
T140D
F
Fetal ISC signature
NES:1.594
FDR: 0.045
WT
T140D
G
YAP signature
NES:1.588
FDR: 0.036
WT
T140D
H
T140D signature up
T140D signature down
Survival probability
p = 0.036
Time

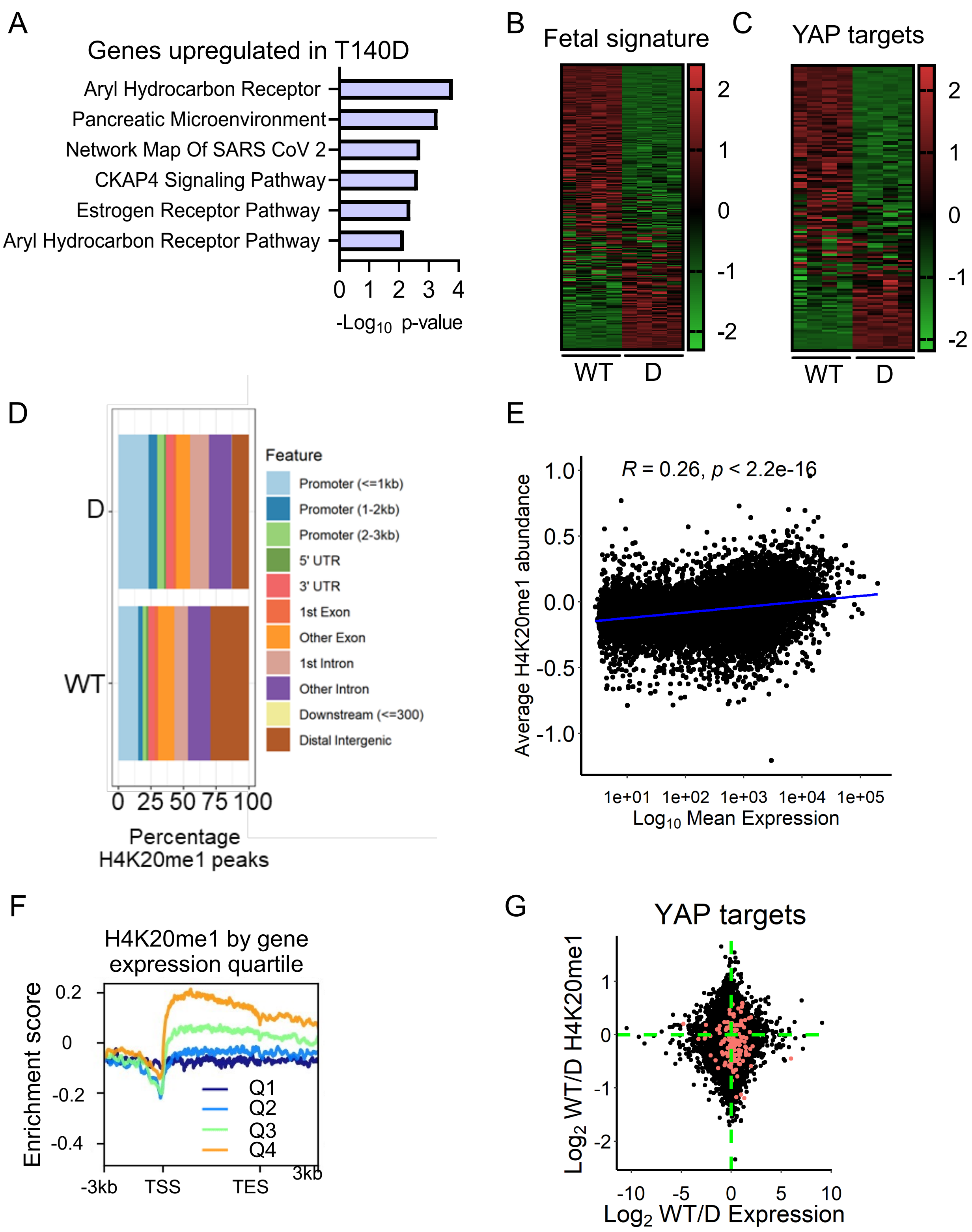


Supplemental Figure 4

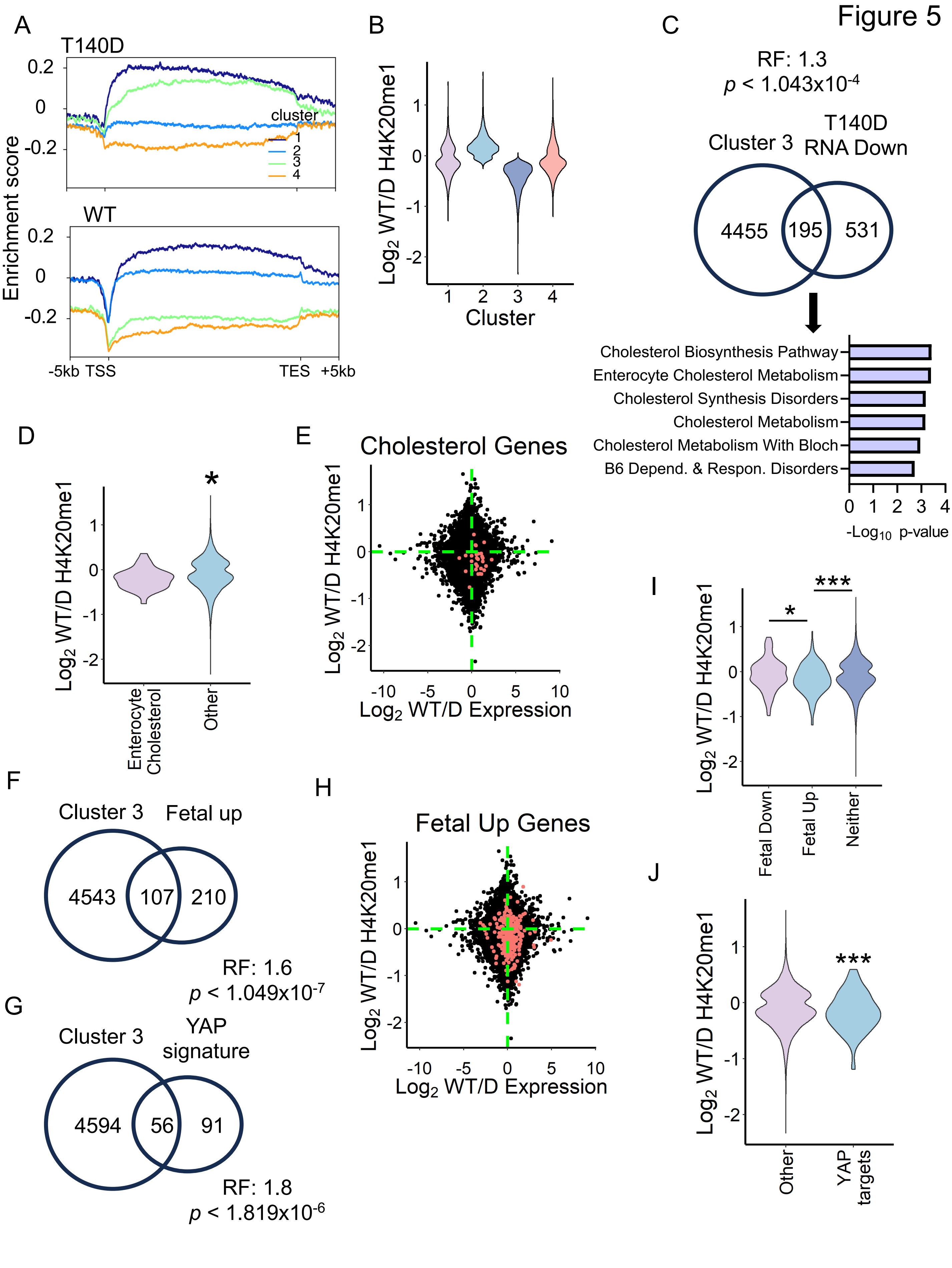

Figure 5
A
T140D
WT
Enrichment score
cluster
1
2
3
4
-5kb TSS
TES +5kb
B
Log2 WT/D H4K20me1
Cluster
C
RF: 1.3
p < 1.043x10-4
Cluster 3
T140D RNA Down
4455
195
531
Cholesterol Biosynthesis Pathway
Enterocyte Cholesterol Metabolism
Cholesterol Synthesis Disorders
Cholesterol Metabolism
Cholesterol Metabolism With Bloch
B6 Depend. & Respon. Disorders
-Log10 p-value
D
Enterocyte Cholesterol
Other
E
Cholesterol Genes
Log2 WT/D Expression
F
Cluster 3
Fetal up
4543
107
210
RF: 1.6
p < 1.049x10-7
G
Cluster 3
YAP signature
4594
56
91
RF: 1.8
p < 1.819x10-6
H
Fetal Up Genes
I
Fetal Down
Fetal Up
Neither
J
Other
YAP targets

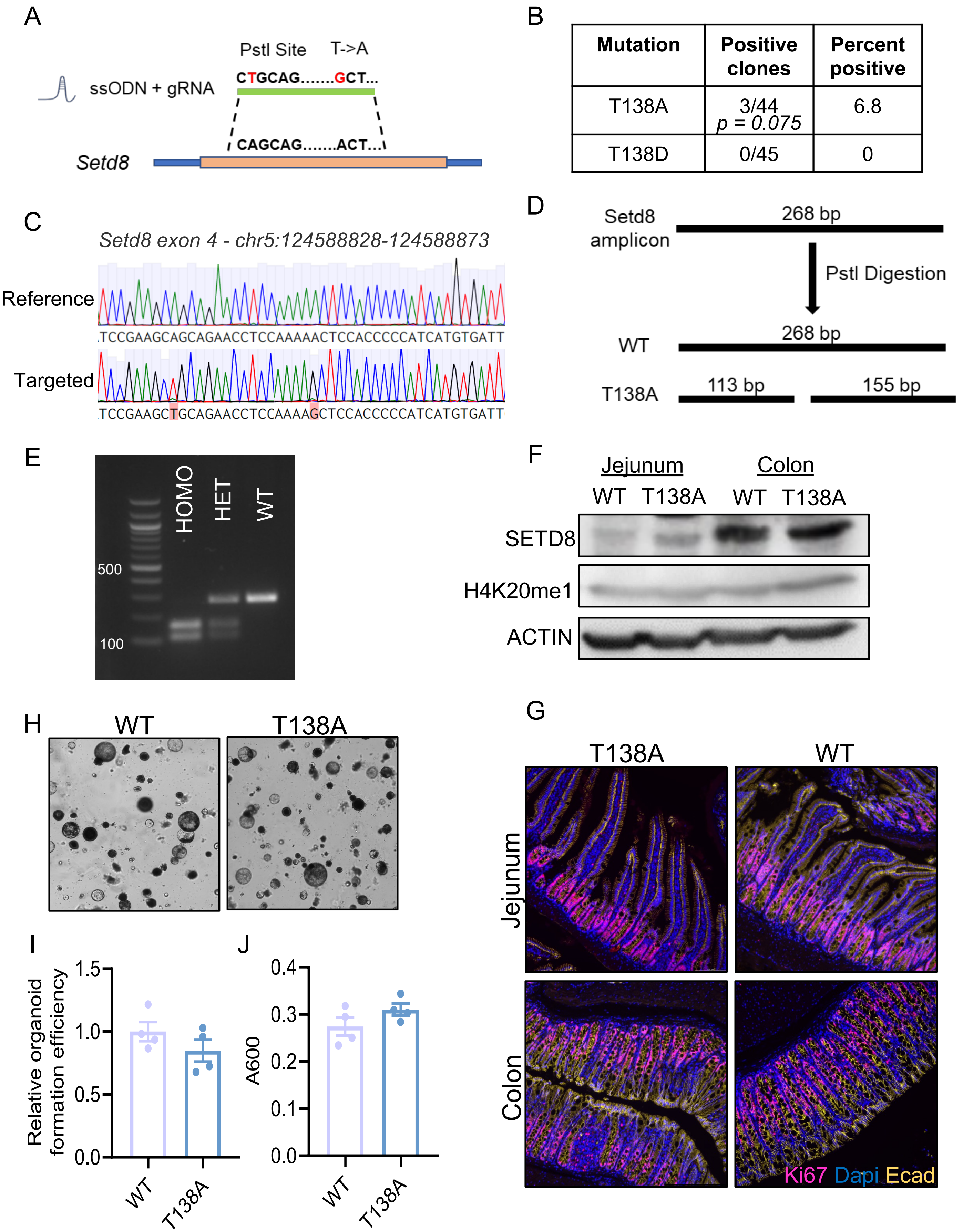


| Mutation | Positive clones | Percent positive |
|---|---|---|
| T138A | 3/44 $p = 0.075$ | 6.8 |
| T138D | 0/45 | 0 |

Supplemental Figure 5

Figure 6

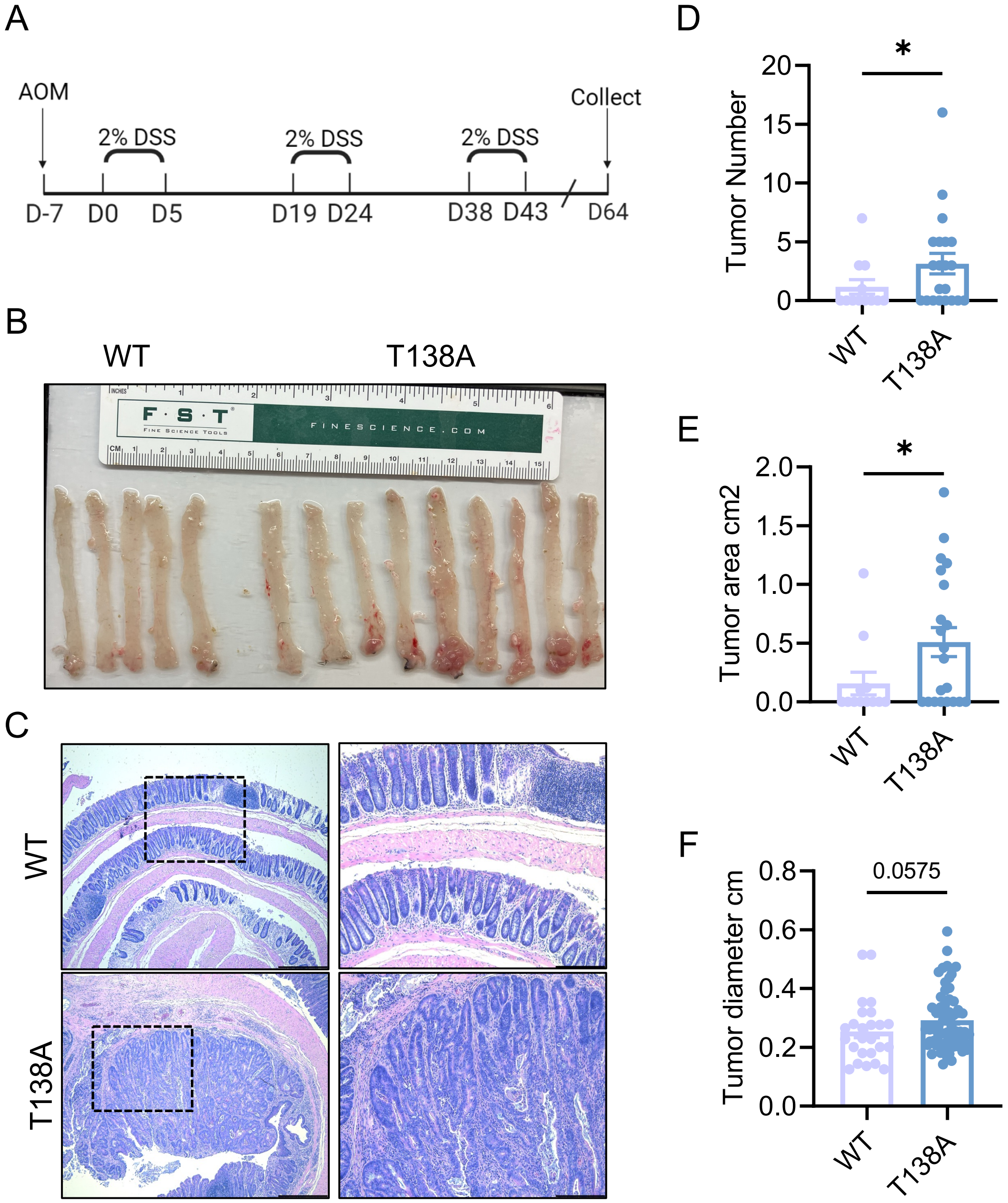
A
AOM
2% DSS
2% DSS
2% DSS
Collect
D-7
D0
D5
D19
D24
D38
D43
D64
B
WT
T138A
C
WT
T138A
D
Tumor Number
*
WT
T138A
E
Tumor area cm2
*
WT
T138A
F
Tumor diameter cm
0.0575
WT
T138A

Fig S1E H4K20me1

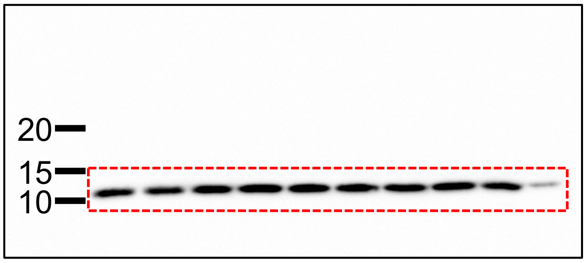


Fig S1E SETD8

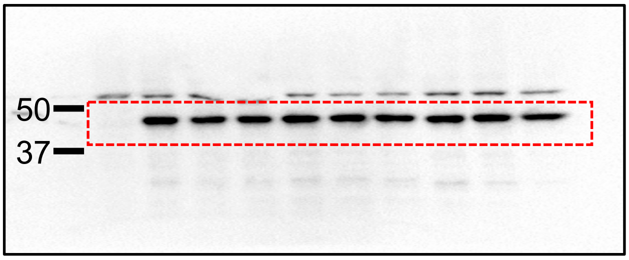


Fig S1E ACTIN

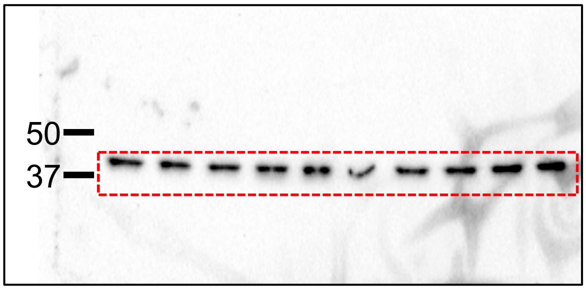


Fig S1E ACTIN

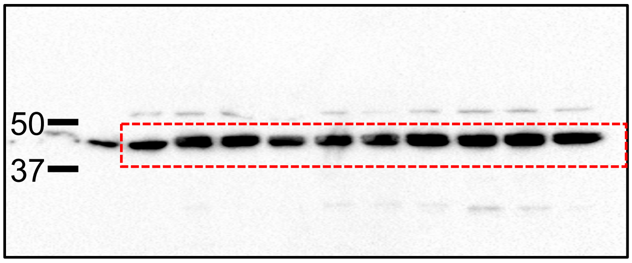


Fig 2D SETD8

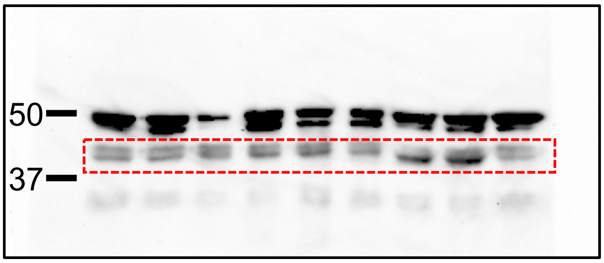


Fig 2D ACTIN

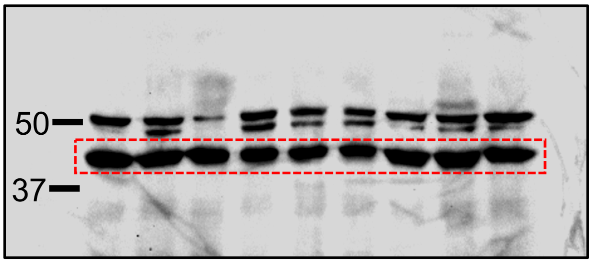


Fig 2D H4K20me1

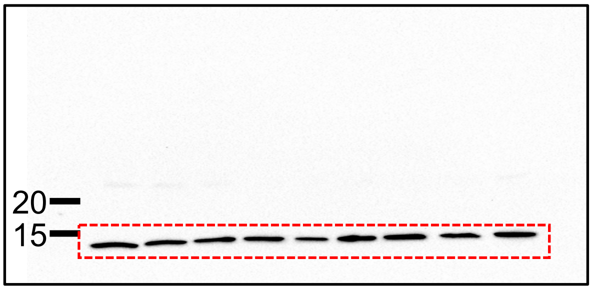


Fig 2D ACTIN

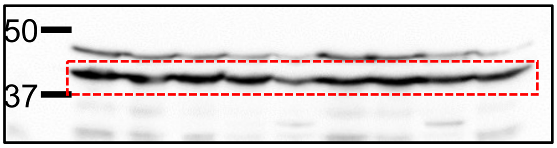

Fig 2I H3 (top) and H4K20me1 (bottom)

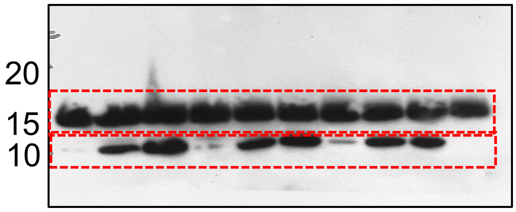


Fig 2E Autoradiograph

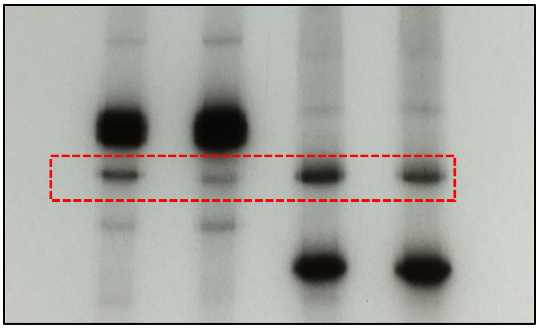

Fig 2J GST-SETD8

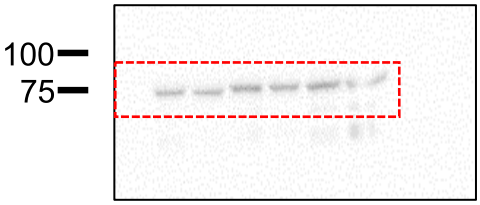


Fig 2E Coomassie

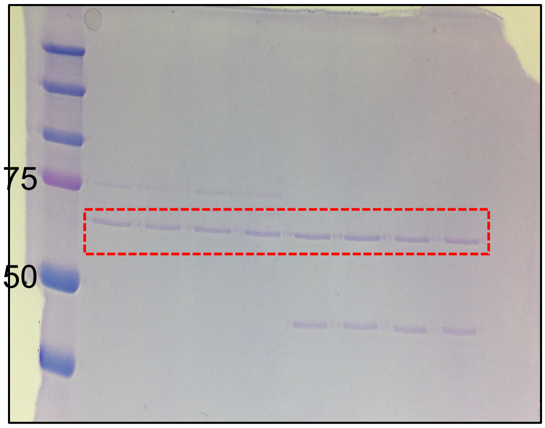


Fig 2J H3

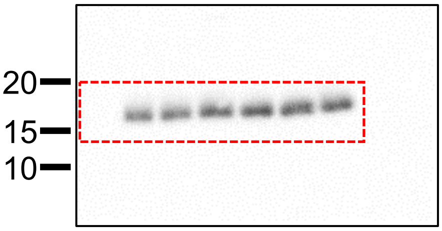


Fig 2F Autoradiograph

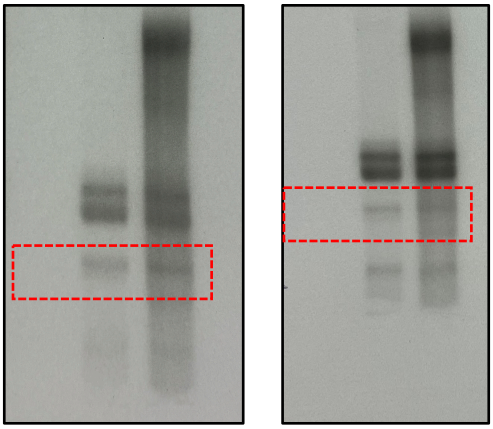

Fig 2F Coomassie

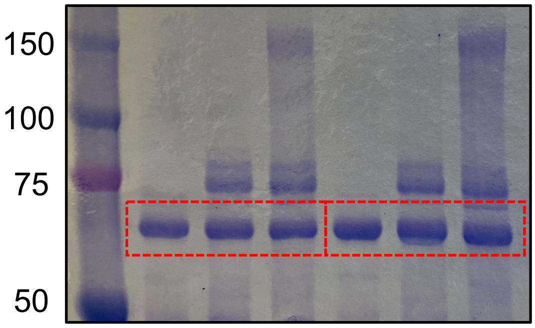

Fig S3B ACTIN

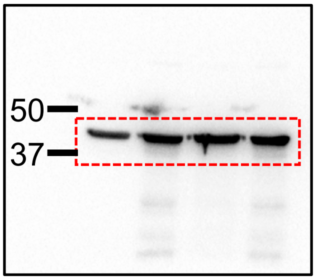


Fig S3C ACTIN

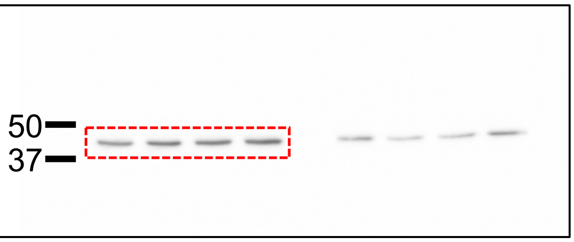


Fig S3C H4K20me1

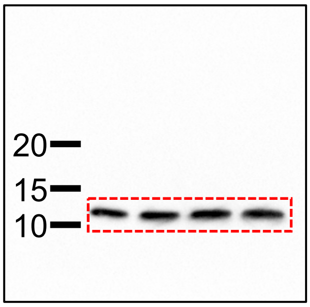


Fig S3B SETD8

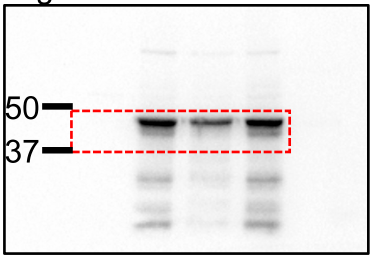


Fig S5F SETD8

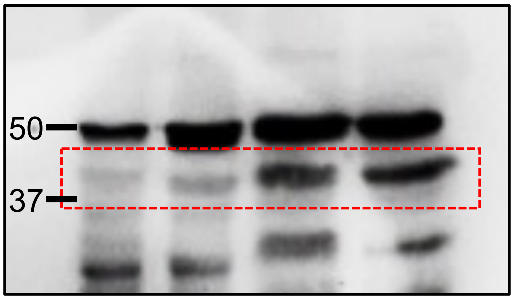


Fig S5F H4K20me1

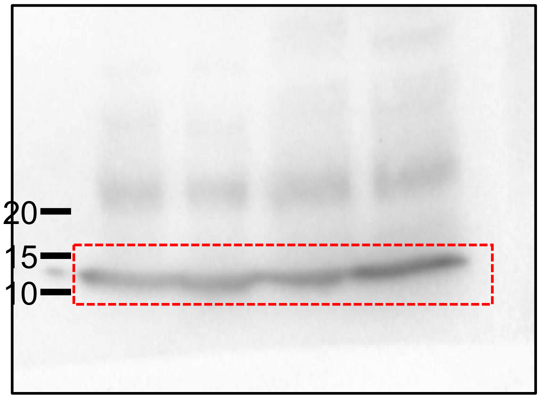


Fig S5F ACTIN

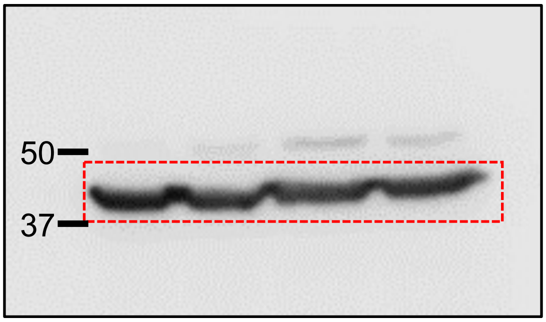
